# Lithosyntrophy: Obligate syntrophy in a phosphite-oxidizing, methanogenic culture

**DOI:** 10.1101/2025.08.26.672485

**Authors:** Heidi S. Aronson, Matt E. Weaver, Ruiwen Hu, V. Celeste Lanclos, Jacob D. Rapp, Anthony T. Iavarone, Hans K. Carlson, John D. Coates

## Abstract

The anaerobic conversion of organic matter to methane and carbon dioxide typically relies on obligate syntrophic interactions between bacteria and methanogenic archaea, where interspecies hydrogen (H_2_) transfer enables thermodynamically constrained reactions to proceed near equilibrium. Syntrophs couple the oxidation of fermentation products such as fatty acids and alcohols to the reduction of protons to form H_2_. These reactions can only proceed if low H_2_ concentrations are maintained by H_2_-consuming syntrophic partners. Here, we describe “lithosyntrophy,” a novel mode of syntrophic interaction in which electrons that drive hydrogenotrophic methanogenesis originate from an inorganic compound rather than from the canonical organic substrates. *Candidatus* Phosphitivorax anaerolimi strain Phox-21 oxidizes phosphite (HPO_3_^2−^, oxidation state +3) to phosphate coupled to hydrogenogenesis in an obligate energetic dependency on a hydrogenotrophic methanogen, *Methanoculleus* sp. Physiology experiments, thermodynamic calculations, genomic annotation, and metaproteomics analysis collectively revealed a mechanism for syntrophic phosphite oxidation in Phox-21, which requires phosphite, acetate, and CO_2_ as co-substrates. In this pathway, electrons derived from phosphite drive H_2_ production via an electron-confurcating hydrogenase. Unlike previously characterized acetogenic phosphite oxidizers that grow without exogenous acetate, Phox-21 requires acetate to regenerate AMP, a cofactor required by the phosphite dehydrogenase, PtdF. Lithosyntrophic phosphite oxidizers may play important roles both in transferring reducing equivalents as well as biologically available phosphorus to other members of their surrounding microbial communities. We infer that lithosyntrophic DPO emerged before acetoclastic methanogenesis and was a major sink for acetate in the Archaean when phosphite was more abundant.

**Significance statement:** Dissimilatory phosphite-oxidizing microorganisms (DPOM) use phosphite as an energy source, producing phosphate. While the two previously isolated DPOM couple phosphite oxidation to carbon fixation via the Wood-Ljungdahl pathway, *Candidatus Phosphitivorax anaerolimi* strain Phox-21 instead performs lithosyntrophic metabolism, coupling DPO to proton reduction in obligate partnership with a hydrogenotrophic methanogen. This establishes a novel link between phosphorus and carbon redox cycles: like organosyntrophy, DPO can fuel methanogenesis. Lithosyntrophy expands our understanding of syntrophic interactions and suggests similar processes may occur with other inorganic substrates in anoxic ecosystems. We propose lithosyntrophic DPO predates acetoclastic methanogenesis, serving as an early acetate sink during the Archaean. Following Earth’s oxygenation, phosphite depletion likely reduced competition for acetate, enabling the later evolution of acetoclastic methanogenesis.

## Introduction

In anoxic environments, the degradation of organic matter to methane and carbon dioxide depends on obligate syntrophic interactions between proton-reducing bacteria and methanogenic archaea (McInerney et al., 2009; Schink, 1997; Stams & Plugge, 2009). Obligate syntrophy is defined by a metabolic and thermodynamic interdependence, where the complete degradation of a substrate requires the activity of two or more microorganisms. Neither syntrophic partner can degrade the organic compound alone, and both catalyze catabolic reactions that proceed close to thermodynamic equilibrium, such that the energetics of one partner’s catabolism is directly influenced by the other. In other words, the accumulation of end products from one organism’s metabolism can thermodynamically inhibit the reaction, and the syntrophic partner drives the reaction forward by removing inhibitory end products. In organotrophic syntrophic partnerships (“organosyntrophy”), syntrophic bacteria couple the oxidation of organic acids and alcohols to the reduction of protons to form hydrogen (H_2_), which is immediately consumed by a hydrogenotrophic methanogen (Bryant et al., 1967; Jackson et al., 1999; McInerney et al., 1979; Zinder & Koch, 1984). The tight metabolic coupling between partner organisms is facilitated by interspecies H_2_ transfer. The drawdown of H_2_ by the methanogen enables the otherwise endergonic (thermodynamically unfavorable, Δ*G*_*r*_ > 0) catabolic reaction of the syntrophic bacterium to become exergonic (thermodynamically favorable, Δ*G*_*r*_ < 0).

While obligate syntrophy has traditionally been associated with the anaerobic degradation of organic substrates (Bryant et al., 1967; McInerney et al., 1979, 2009; Schink, 1997), the underlying thermodynamic principles are not limited to organotrophy. In principle, any oxidation reaction that produces an inhibitory accumulation of H_2_ may require syntrophic coupling to proceed. However, such interactions involving inorganic electron donors have not been previously described. While H_2_ formation from FeS and H_2_S (Wächterhäuser reaction) has been shown to support hydrogenotrophic methanogenesis, the biological mechanism supporting this reaction is unclear (J. Thiel, 2020; J. Thiel et al., 2019), and abiotic H_2_ formation from FeS has been shown to occur (Helmbrecht et al., 2025). Here, we report the discovery of phosphite-driven “lithosyntrophy,” a novel mode of obligate syntrophic interaction in which electrons that drive hydrogenotrophic methanogenesis originate from an inorganic compound rather than from an organic substrate.

This discovery builds on prior studies of dissimilatory phosphite oxidation (DPO), a microbial catabolism by which phosphite (HPO_3_^2-^, oxidation state +3) is oxidized to phosphate. Since the first demonstration of DPO in 2000 (Schink & Friedrich, 2000), only two DPO microorganisms (DPOM) have been isolated in pure culture. *Desulfotignum phosphitoxidans* FiPS-3 and *Phosphitispora fastidiosa* DYL19 were isolated from enrichments that used phosphite as the sole electron donor, CO_2_ as the sole carbon source, and sulfide as a reducing agent (Mao et al., 2021; Schink et al., 2002). In these enrichments, CO_2_ also served as the electron acceptor for DYL19, while sulfate was included as the electron acceptor for FiPS-3. Both isolates are acetogens that couple phosphite oxidation to carbon fixation (Table 1, Rxn. 1) through the Wood-Ljungdahl pathway, which is the only known carbon fixation pathway that is coupled to energy conservation (Schuchmann & Müller, 2014), while FiPS-3 can also couple phosphite oxidation to dissimilatory sulfate reduction. In these organisms, acetate is produced as a detectable byproduct (Mao et al., 2021; Mao, Müller, et al., 2023).

*Candidatus* Phosphitivorax anaerolimi Phox-21, the third described DPOM, was enriched from wastewater treatment sludge in chemically defined media that contained phosphite as the primary electron donor, CO_2_/bicarbonate as the exogenous electron acceptor and cysteine as a reducing agent (Ewens et al., 2021; Figueroa et al., 2017). Based on its genomic potential for the reductive glycine pathway, and its absolute dependency on CO_2_ for growth, it was initially assumed that, like FiPS-3 and DYL19, Phox-21 was an autotroph coupling DPO to carbon fixation (Ewens et al., 2021; Figueroa et al., 2017). However, several lines of evidence suggested that Phox-21 is physiologically distinct from FiPS-3 and DYL19. Phox-21 was resistant to isolation via dilution-to-extinction or colony picking from agar plugs, which is consistent with its phylogenetic relatedness to fastidious syntrophic bacteria within the phylum Desulfobacterota (Ewens et al., 2021; Figueroa et al., 2017). In contrast to FiPS-3, Phox-21 lacks an electron transport chain and pathways for alternative metabolic strategies such as anaerobic respiration or fermentation, which are characteristics of metabolically specialized syntrophs (McInerney et al., 2009). Furthermore, Phox-21 lacks genes for phosphoacetyltransferase (PTA) and acetate kinase (ACK) within the reductive glycine pathway (Ewens et al., 2021), preventing acetate production and ATP generation via substrate-level phosphorylation (Sánchez-Andrea et al., 2020). Without energy conservation from the reductive glycine pathway, it was unclear what the terminal electron acceptor for phosphite was. Collectively, these genomic, metabolic, and phylogenetic features suggested that Phox-21 is physiologically distinct from isolated DPOM and depends on syntrophic interactions with other microorganisms to oxidize phosphite.

In this study, we describe lithosyntrophic phosphite oxidation, an uncharacterized mode of obligate syntrophy in which Phox-21 couples the oxidation of phosphite to proton reduction, generating H_2_ that is consumed by a hydrogenotrophic methanogen, *Methanoculleus* sp. By demonstrating that inorganic compounds can sustain methanogenic ecosystems, these findings establish a mechanistic link between the phosphorus and carbon redox cycles and challenge the assumption that obligate syntrophy is inherently linked to organic carbon degradation (McInerney et al., 2009; Schink, 1997). Lithosyntrophy may represent a historically significant and previously overlooked contributor to methanogenesis, and similar interactions may be more widespread in contemporary anoxic environments than previously appreciated. This work provides new insights into the thermodynamic and ecological constraints that shape microbial community structure and function in methanogenic environments and raises new questions about the environmental distribution and evolutionary origins of phosphorus-based energy metabolisms.

## Results

### Electrons from phosphite are stoichiometrically balanced with methanogenesis or sulfate reduction

As of July 2024, highly enriched phosphite-oxidizing cultures maintained with cysteine as a reducing agent supported a stable microbial community comprising 15 taxa, including *Candidatus* Phosphitivorax anaerolimi Phox-21, *Methanoculleus sp., Acetomicrobium hydrogeniformans*, and other fermentative bacteria (Fig. 1A). Removing CO_2_ or cysteine from the medium arrested growth and phosphite oxidation (Ewens, 2022; Ewens et al., 2021). Sulfide alone could not replace cysteine as a reducing agent, which suggested cysteine, or a degradation product of cysteine, was necessary for phosphite oxidation to occur. Based on the physiology of their closest isolated relatives, we hypothesized that *Acetomicrobium, Aminobacterium, and Aminivibrio* were likely fermenting cysteine to produce sulfide, ammonia, and pyruvate (Hamdi et al., 2015; Honda et al., 2013; Maune & Tanner, 2012; Yokota & Ikeda, 2017). Because organic acids like pyruvate are fermentatively catabolized to acetate, we chose to supplement the media with acetate, a non-fermentable carbon source, rather than pyruvate, to simplify the organic input into the system. By replacing cysteine with acetate (3 mM) and sulfide (1 mM), the community composition was reduced to 7 members with the following GTDB taxonomy: Phox-21, *Methanoculleus* sp018433835*, Acetomicrobium hydrogeniformans, Desulfomicrobium apsheronum, Proteiniphilum massiliensis, Mesotoga prima,* and an unclassified member of the class Peptococcia (order DRI-13, genus SV3-B141), with Phox-21 comprising >98% of the community (Fig. 1B). Attempts to isolate Phox-21 on acetoin or crotonate instead enriched for the unclassified strain of Peptococcia (Supp. Fig. 1)

**Figure 1:**
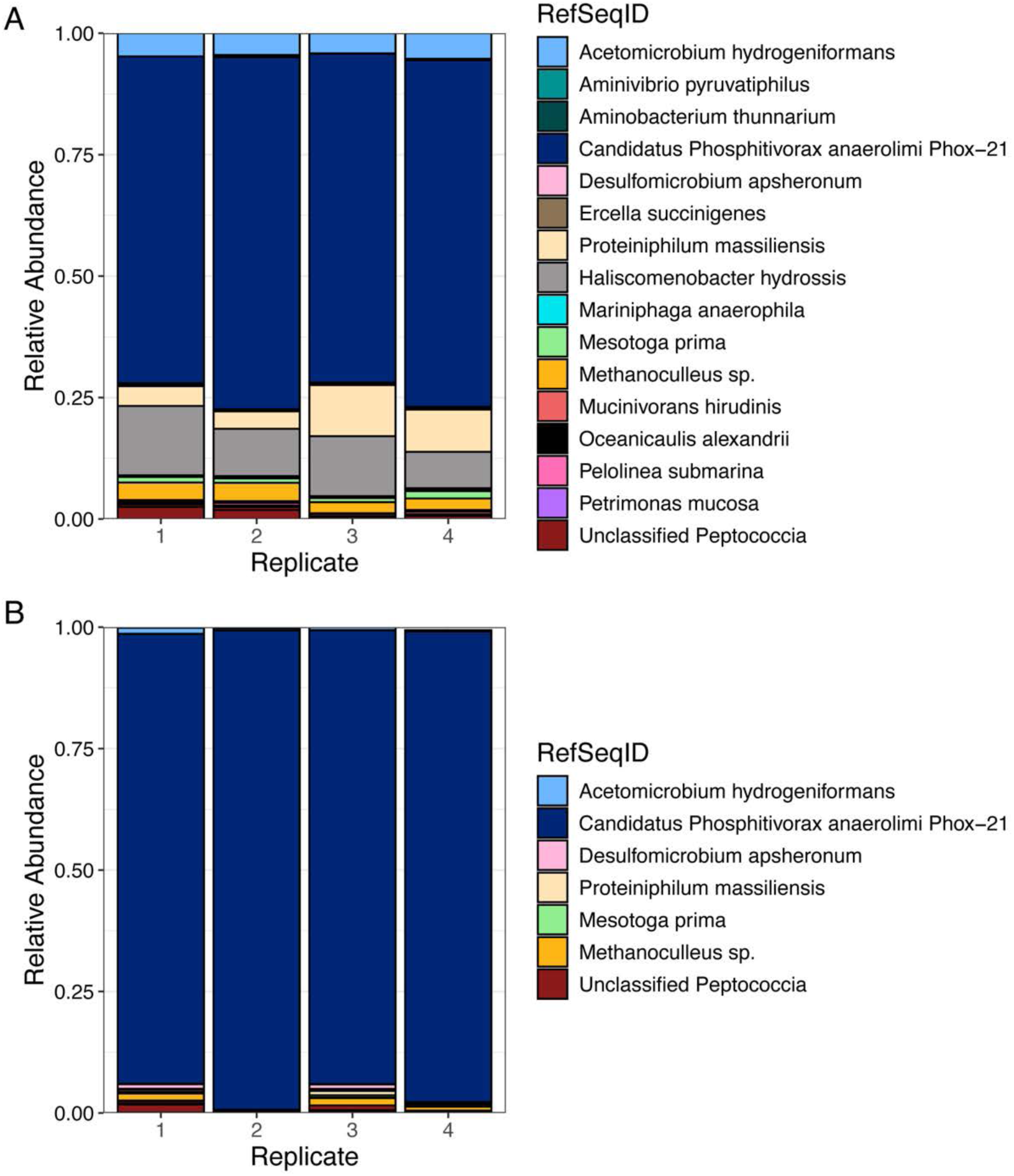
Relative abundances of microbial taxa from phosphite-oxidizing enrichment cultures A) with cysteine and B) with acetate and sulfide. Each bar represents a replicate culture. Taxonomic identification is based on the V4/V5 16S rRNA gene region.

In the sulfide-reduced cultures with acetate, growth reached maximum optical density with concurrent complete oxidation of 10 mM phosphite to 10 mM phosphate after 4 days of incubation (Fig. 2). Aqueous methane concentration reached a maximum of ∼30 µM (0.032 atm CH_4(g)_). Phosphite oxidation was stoichiometric with methane production in the methanogenic phosphite-oxidizing culture (Table 1, Rxn. 4; Fig. 3A). In the presence of the methanogenic inhibitor 2-bromoethanesulfonate (BES, 1 mM), growth and phosphite oxidation of the methanogenic phosphite oxidizing culture were inhibited (Fig. 2)

**Figure 2:**
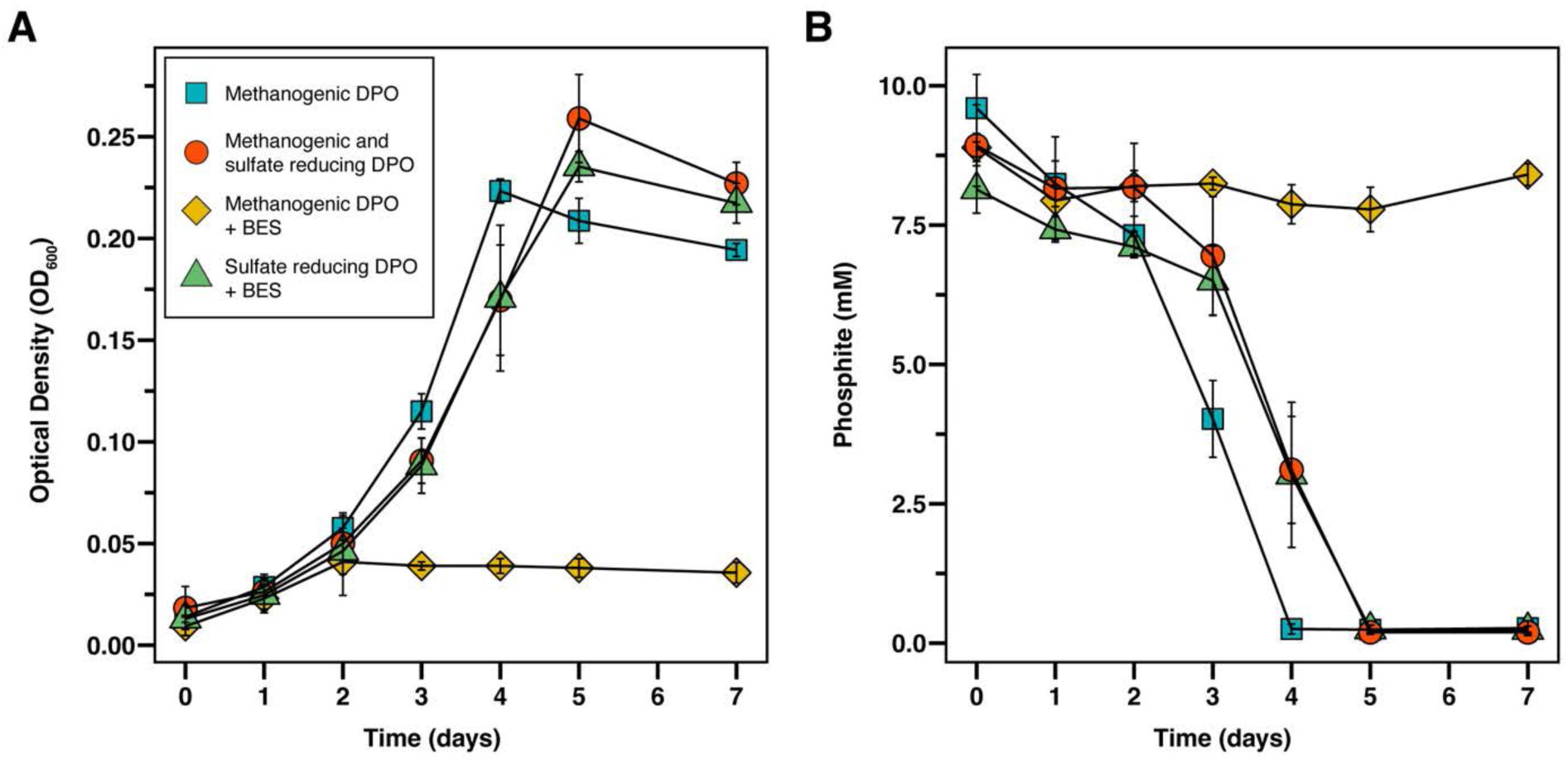
Growth (A) and phosphite concentration (B) of phosphite-oxidizing enrichment cultures under methanogenic conditions (blue squares), with 1 mM BES (yellow diamonds), methanogenic and sulfate-reducing conditions with *D. desulfuricans* G11 (red circles), and sulfate-reducing conditions with *D. desulfuricans* G11 and 1 mM BES (green triangles). Error bars represent SD of triplicate cultures.

**Figure 3:**
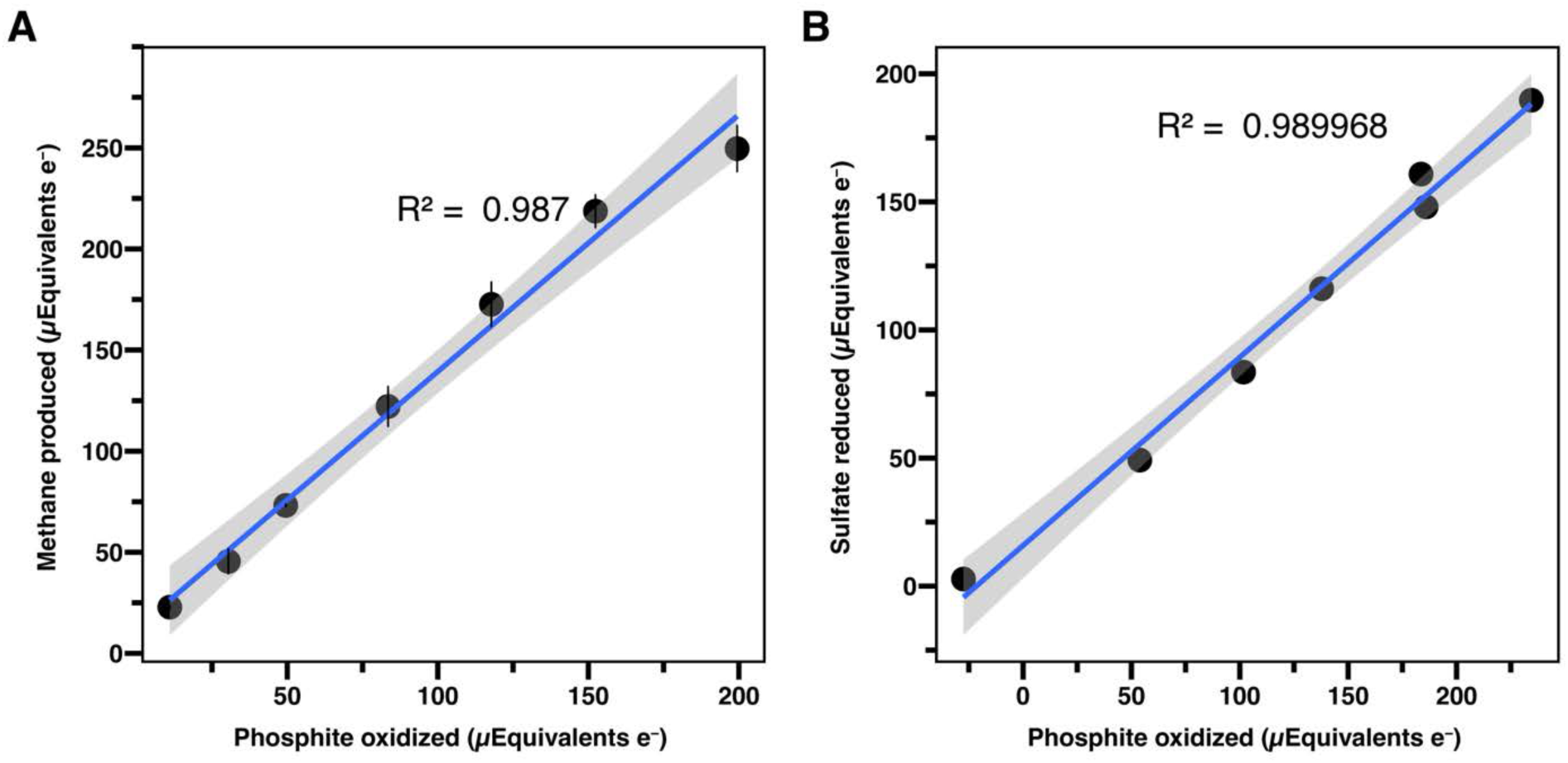
Relationships between phosphite oxidation and terminal electron-accepting processes under methanogenic conditions (A) or sulfate-reducing conditions (B). Concentrations of phosphite, methane, and sulfate were normalized by µM electron equivalents.

### Acetate is required but is not the source of methane

A minimum concentration of 800 µM acetate was required for growth and phosphite oxidation to occur (Supp. Figs. 2 and 3). Because of the long-term maintenance of a methanogen, *Methanoculleus* sp., in the enrichment, we considered the possibility that acetate was required for acetoclastic methanogenesis. Measurements of methane via GC-MS in cultures incubated with ^13^C bicarbonate or ^13^C acetate showed that carbon from acetate was not the source of methane (Supp. Fig. 4). Methane measured in the condition incubated with ^13^C bicarbonate had a significantly higher enrichment of ^13^C/^12^C compared to the unlabeled control (p = 2.31e-13), while the conditions incubated with ^13^C acetate produced methane that was not significantly enriched compared to the unlabeled control (p = 0.195 and 0.386). This was supported by physiological analysis of the *Methanoculleus* sp. isolate, which could only produce methane from H_2_/CO_2_ or formate, but not acetate (Supp. Fig. 5).

To better understand Phox-21’s carbon metabolism and its requirement for acetate, incubations were amended with ^13^C-labeled bicarbonate or ^13^C-labeled acetate. Based on analysis of the mass spectra from abundant peptides detected by metaproteomics, carbon from bicarbonate and acetate was incorporated into peptides produced by Phox-21. Within the peptide YDLVEGLK (z = 2^+^), which matched to the protein sequence of the NAD^+^/AMP-dependent phosphite dehydrogenase PtdF/ApdA, there was one carbon substitution from ^13^C bicarbonate and 15 carbon substitutions from ^13^C acetate, (Supp Fig. 6), indicating that Phox-21 assimilates carbon from both bicarbonate and acetate. This is consistent with the presence and activity of the carbon fixing reductive glycine pathway in Phox-21’s genome, the presence of these proteins in the metaproteome, and the physiological requirement for acetate (Supp. Figs. 2, 3, and 4). The in-depth carbon metabolism of Phox-21 and the relative contributions of autotrophy vs. heterotrophy is the subject of current investigations and will be described in a separate publication.

### DPO requires and maintains low H_2_ concentrations

If Phox-21 is a syntroph, its metabolism likely depends on interspecies electron transfer to the partner organism, *Methanoculleus* sp. Because *Methanoculleus* sp. produces methane only from H_2_/CO_2_ and formate (Supp. Figs. 3 and 4), we hypothesized that Phox-21 couples phosphite oxidation to proton reduction to produce H_2_ (Table 1, Rxns. 4 and 5). This is consistent with interspecies H_2_ transfer, a common form of electron exchange in organosyntrophic interactions (Stams & Plugge, 2009). Hydrogenogenesis from proton reduction is energetically difficult because of the low midpoint redox potential of the H^+^/H_2_ couple (*E*^0′^= −414 mV; Δ*G*^0^ = +18.6 kJ/mol) and is thus sensitive to the accumulation of H_2_. The syntrophic partner organism continuously removes and maintains low H_2_ concentrations that are thermodynamically necessary for the reaction to proceed. This mechanism is widely observed in organosyntrophic interactions involving methanogens and provided a plausible framework for understanding Phox-21’s catabolism and its relationship with *Methanoculleus* sp.

To confirm that Phox-21 requires a H_2_ -consuming partner, we inoculated the culture with the model hydrogenotrophic sulfate reducer *Desulfovibrio desulfuricans* G11 in the presence 1 mM BES (Fig. 2). Growth and phosphite oxidation were restored, and stoichiometric phosphite oxidation and sulfate consumption was observed under sulfate-reducing conditions, which was consistent with the syntrophic coupling of phosphite oxidation to hydrogenotrophic sulfate reduction (Table 1, Rxn. 6; Fig 3B).

Threshold H_2_ activities for thermodynamic favorability were calculated from the Gibbs energy of phosphite oxidation coupled to proton reduction (Rxn. 5), hydrogenotrophic methanogenesis (Rxn. 7), hydrogenotrophic sulfate reduction (Rxn. 8), and phosphite oxidation coupled to proton reduction, NAD^+^ reduction, and ADP formation (Rxn. 9) as a function of H_2(aq)_ activity (Fig. 4). Under in vitro conditions of the enrichment culture, calculations predict that hydrogenotrophic methanogenesis requires a minimum hydrogen activity of −8.95 (≈1.12 nM H_2(aq)_). Lithosyntrophic phosphite oxidation coupled to sulfate reduction (Rxn. 6) has a lower threshold H_2_ concentration (0.275 nM) than methanogenic phosphite oxidation, which is consistent with the observation that sulfate reducers have a lower K_s_ for H_2_ than sulfate reducers do (Fig. 4) (Kristjansson et al., 1982).

**Figure 4:**
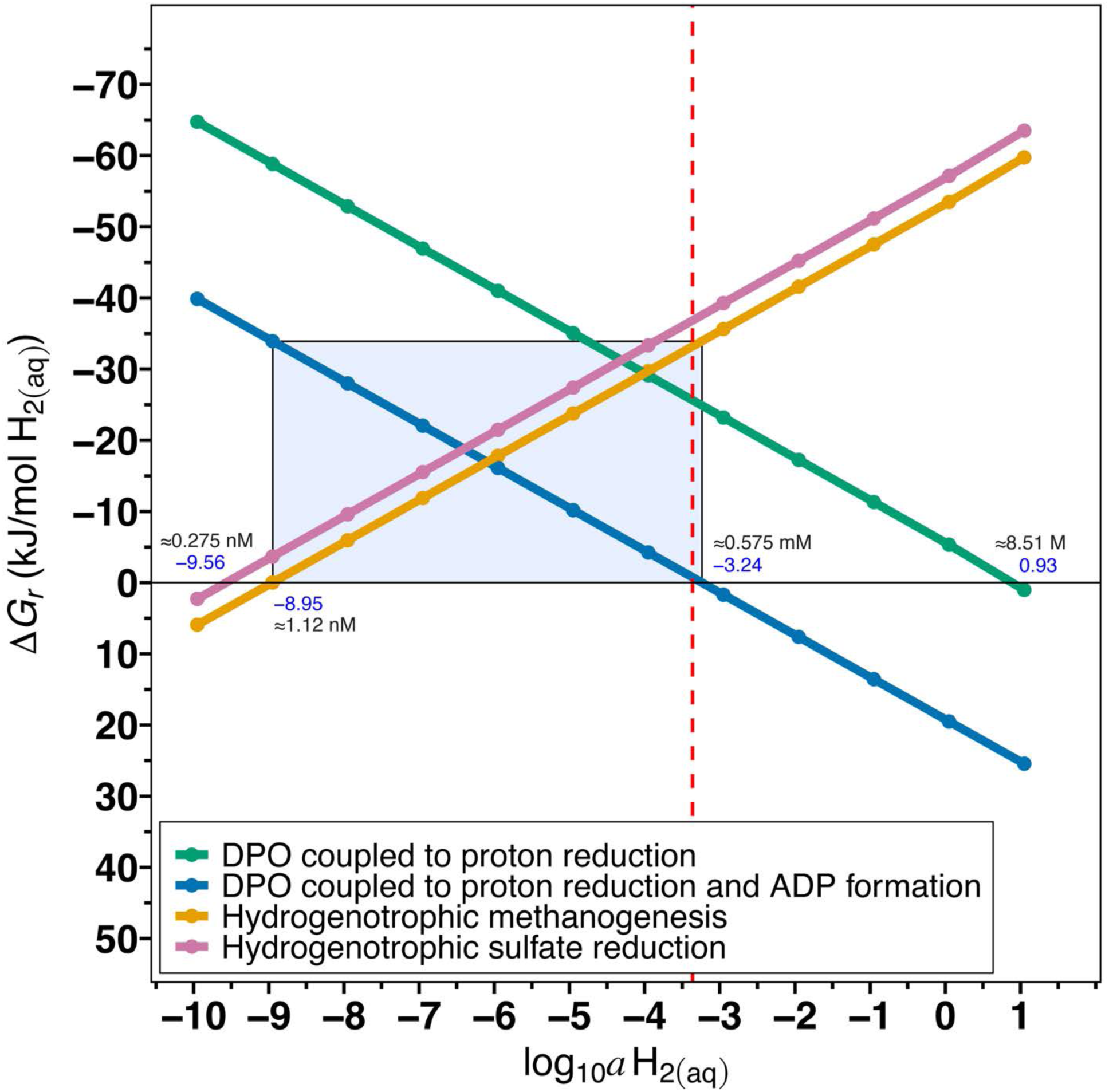
Gibbs energies, Δ*G_r_*, of reactions 10, 11, 12, and 16 (Table 1) as a function of hydrogen activity. Activities (Supp. Table X) for products and reactants were calculated from the concentrations listed in Supp. Table X. The blue box indicates the range of hydrogen activities where hydrogenotrophic methanogenesis and DPO coupled to proton reduction and ADP formation are both exergonic. The red dashed line indicates the aqueous hydrogen activity equivalent to adding 10 mL of hydrogen gas to the headspace of a culture tube. *ΔG_r_* was calculated using activities determined from the following concentrations: 10 mM phosphite, 100 µM methane, 10 mM sulfate, 2.2 mM ATP, 0.83 mM ADP, 0.3 mM AMP, and 100 µM NAD^+^ and NADH at 37 °C.

The energy yield of phosphite oxidation coupled to proton reduction (Table 1, Rxn. 5) is uninhibited by aqueous H_2_ activities below 0.93 (∼8.51 M H_2(aq)_) (Fig. 4) and theoretically should not require hydrogen removal to remain exergonic. This reaction yields −52.9 kJ mol^−1^ H_2_ under the *in vitro* H_2_ concentrations measured in the culture (8.2 ± 0.7 nM H_2(aq)_; Supp. Fig. 7). Although the net reaction of phosphite oxidation coupled to proton reduction is exergonic at the measured H_2_ concentrations, growth and phosphite oxidation were unexpectedly inhibited when excess H_2_ was added to the headspace of the tubes (∼0.433 mM H_2(aq)_; Fig. 5). We hypothesized that specific intracellular reactions, particularly those involving H_2_ production from biochemical reducing equivalent carriers (NADH), were thermodynamically constrained by H_2_ accumulation, which led to inhibition of growth and phosphite oxidation.

**Figure 5:**
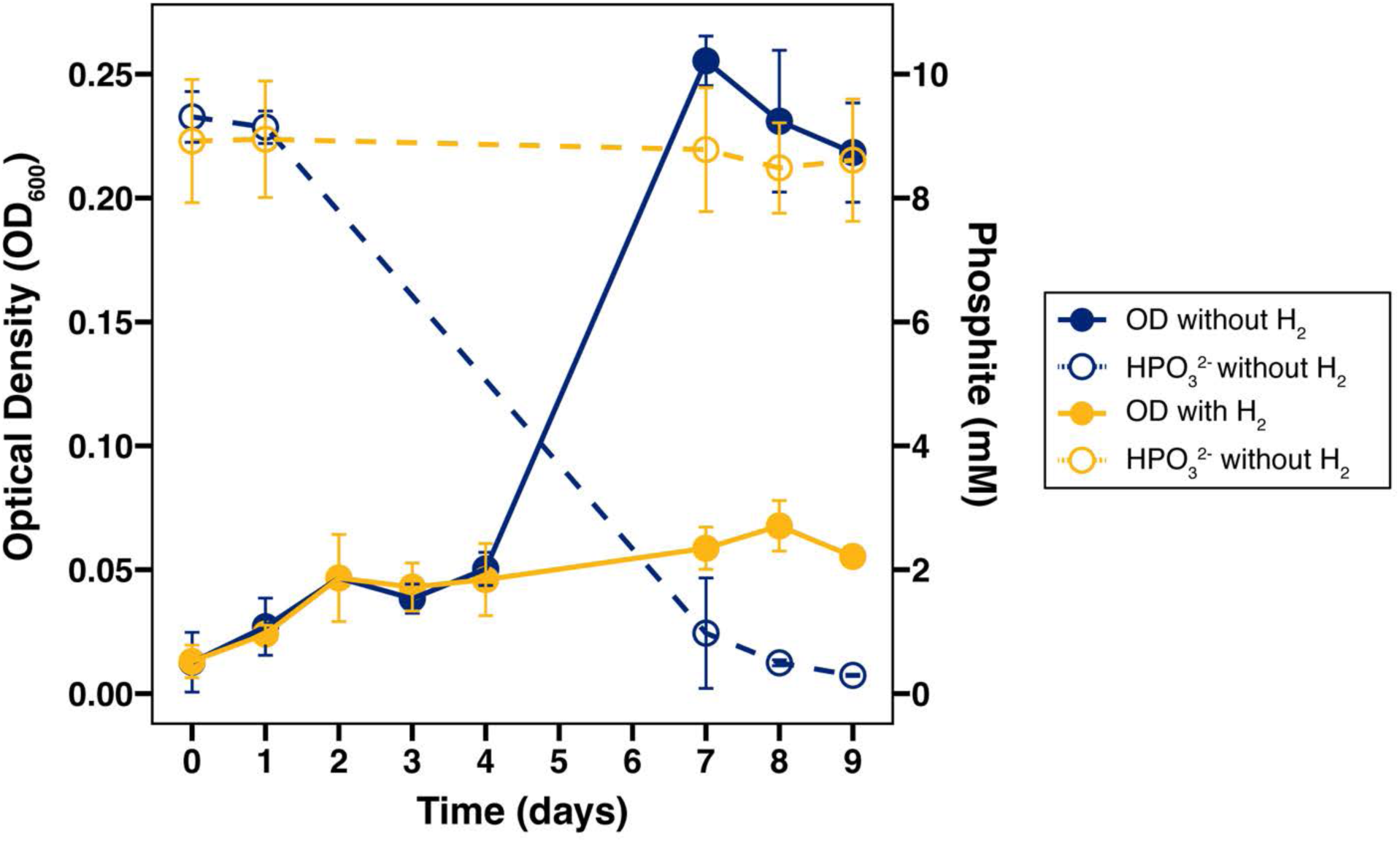
Growth (solid points and lines) and phosphite concentrations (open points, dashed lines) in the presence and absence of 10 mL H_2_ added to the headspace of the culture tubes. Error bars represent SD of triplicate cultures.

### A model of lithosyntrophic phosphite oxidation in Phox-21

To investigate the metabolic basis for this thermodynamic inhibition, we constructed a model of Phox-21’s catabolism using the most abundant peptides detected by metaproteomics (Fig. 6; Supp. Table 4). The most abundant protein was the NAD^+^/AMP-dependent phosphite dehydrogenase, PtdF/ApdA, which is part of the ptx-ptd gene cluster conserved in all DPOM (Ewens et al., 2021; Figueroa et al., 2017; Mao, Fleming, et al., 2023). Other abundant proteins included the putative phosphite-phosphate antiporter PtdC, an NADH- and ferredoxin-oxidizing, electron confurcating/bifurcating hydrogenase HndABCD, a membrane-bound, Na^+^-translocating ferredoxin:NAD^+^ oxidoreductase Rnf complex, an adenylate kinase, acetyl-coenzyme A synthetases, a K^+^-stimulated pyrophosphate-energized Na^+^/H^+^ pump, and ATP synthases. Note that HndC was annotated as a homolog of NADH:quinone oxidoreductase (NuoF) (Kpebe et al., 2018). The Rnf complex, electron-confurcating hydrogenase, and ATP formation through substrate-level phosphorylation are energy conservation mechanisms that are involved in syntrophic metabolism that are central to our proposed model of phosphite oxidation in Phox-21 (Fig. 6) (James et al., 2016; McInerney et al., 2009; J. R. Sieber et al., 2012). Other abundant proteins included several proteins involved in the reductive glycine pathway, including formate dehydrogenase, methylene-THF dehydrogenase, glycine dehydrogenase, serine hydroxymethyltransferase, and serine dehydratase.

**Figure 6:**
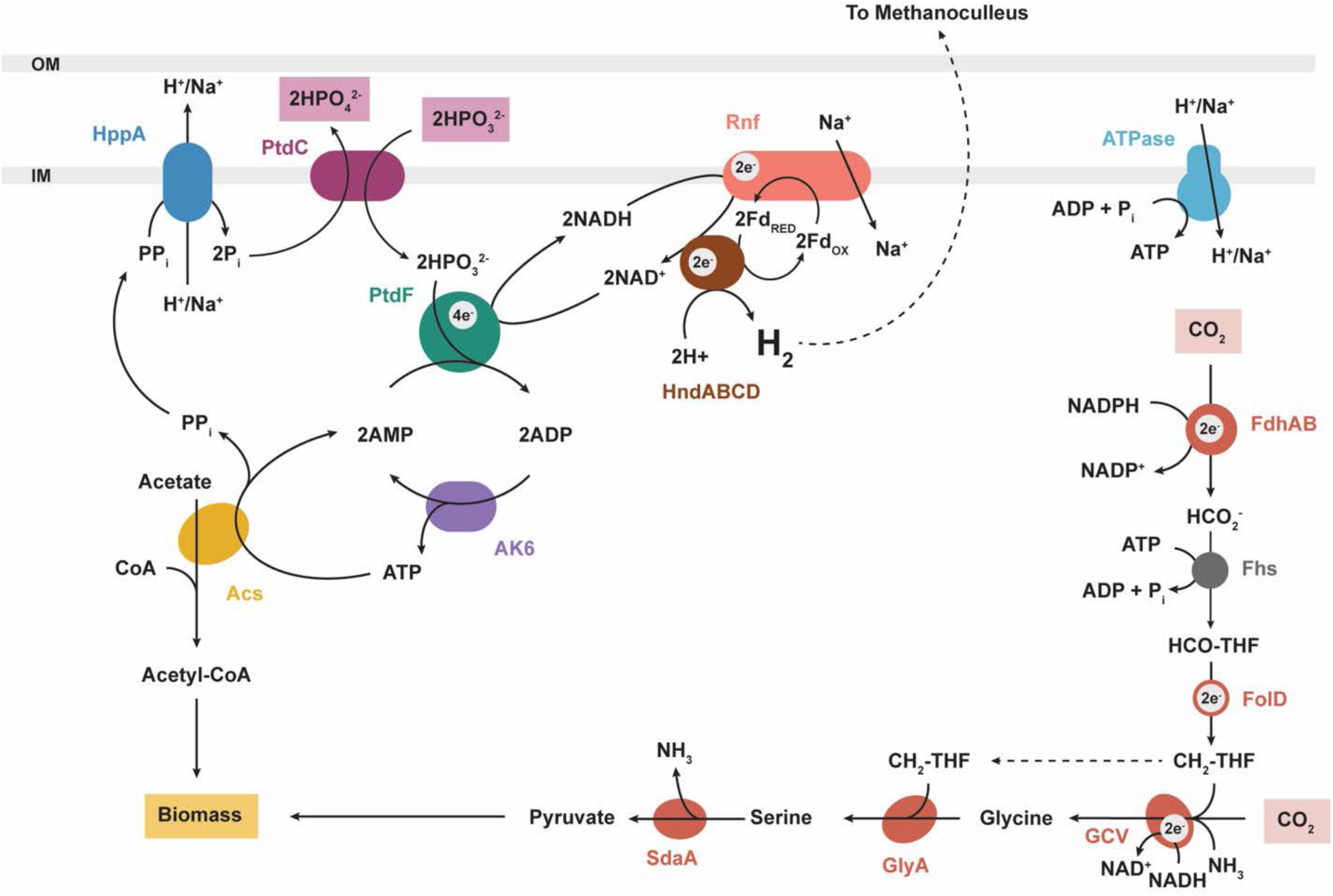
Proteomics-based metabolic model of dissimilatory phosphite oxidation coupled to proton reduction in *Ca.* Phosphitivorax anaerolimi Phox-21. PtdC: phosphite-phosphate antiporter, PtdF: AMP-dependent phosphite dehydrogenase, HndABCD: NADH- and ferredoxin-oxidizing, electron-confurcating hydrogenase, Rnf: membrane-bound, Na^+^-translocating ferredoxin:NAD^+^ oxidoreductase, AK6: adenylate kinase, Acs: acetyl-coenzyme A synthetase, HppA: K^+^-stimulated pyrophosphate-energized Na^+^ pump. Reductive glycine pathway: FdhAB: formate dehydrogenase, Fhs: formate:tetrahydrofolate (THF) ligase, FolD: methylene-THF dehydrogenase/methenyl-THF cyclohydrolase, GCV: glycine cleavage system, GlyA: serine hydroxymethyltransferase, SdaA: serine dehydratase/threonine dehydratase. While Fhs (gray circle) was annotated in the genome, peptides associated with this protein were not detected in the metaproteome.

In our proposed model of lithosyntrophic phosphite oxidation (Fig. 6), phosphite first enters the cell via PtdC, a phosphite-phosphate antiporter. Electrons from phosphite are transferred by PtdF/ApdA to NAD^+^ to generate NADH and phosphate, with concomitant AMP phosphorylation to ADP (Mao, Fleming, et al., 2023). This reaction runs twice (Table 1, Rxn. 10). HndABCD, a cytoplasmic FeFe electron-confurcating hydrogenase, couples the oxidation of reduced ferredoxin and NADH to proton reduction to generate H_2_ (Table 1, Rxn. 11) (Kpebe et al., 2018; Payne et al., 2022; Schut & Adams, 2009). Reduced ferredoxin is produced by the Rnf complex (Table 1, Rxn. 12) at the expense of NADH and a Na^+^ motive force (Biegel et al., 2011; Westphal et al., 2018). Adenylate kinase converts 2ADP to ATP + AMP (Table 1, Rxn. 13). Acetyl-coA synthetase (Acs) converts acetate, ATP, and coA to AMP, pyrophosphate, and acetyl-coA, which is assimilated into biomass (Table 1, Rxn. 14). The ion-translocating pyrophosphate hydrolyzes pyrophosphate into two molecules of inorganic phosphate (Table 1, Rxn. 15), coupling this exergonic reaction to the translocation of protons or sodium ions across the membrane to maintain a proton/sodium motive force. Two molecules of inorganic phosphate are transported out of the cell by PtdC.

The energy yield of reducing protons with NADH to form H_2_ (Table 1, Rxn. 16) is close to thermodynamic equilibrium and is thus very sensitive to the accumulation of H_2_ (Δ*G*^0^ = −6 kJ/mol, Δ*G*_16_ = −9.6 kJ/mol H_2_ at 10 nM H_2(aq)_). This thermodynamic barrier is overcome in organosyntrophic bacteria by coupling the exergonic oxidation of ferredoxin to the endergonic oxidation of NADH to generate H_2_ in an electron confurcation reaction (Table 1, Rxn. 11) (Schut & Adams, 2009; J. R. Sieber et al., 2012). Reduced ferredoxin is generated by the Rnf complex in an energy consuming reaction (Table 1, Rxn. 12; Δ*G*^0^ = +129.4 kJ/mol, Δ*G*_12_ = +132.6 kJ/mol) at the expense of NADH and a Na^+^ motive force (Biegel et al., 2011; Westphal et al., 2018). Using electrons donated from ferredoxin and NADH, HndABCD reduces protons to form H_2_. Under Phox-21’s intracellular conditions, this reaction (Table 1, Rxn. 11) is exergonic (Δ*G*^0^ = −104.2 kJ/mol, Δ*G*_11_ = −37.1 kJ/mol H_2_ at 10 nM H_2(aq)_). The key catabolic reaction of this model catalyzed by PtdF/ApdA, HndABCD, and the Rnf complex, which together link phosphite oxidation to H_2_ production and energy conservation via substrate-level phosphorylation (Table 1, Rxn. 9; Fig 6), yields −28.1 kJ/mol H_2_ and becomes endergonic at an aqueous H_2_ activity −3.24 (≈0.575 mM H_2(aq)_; Fig. 4). Thus, while the net reaction of phosphite oxidation coupled to proton reduction is exergonic, the sensitivity of the hydrogenase to H_2_ concentrations presents a thermodynamic bottleneck that explains Phox-21’s requirement for a hydrogen consuming syntrophic partner. For both phosphite oxidation and hydrogenotrophic methanogenesis to remain exergonic, H_2_ concentrations in the cultures must be maintained at a minimum of 1.12 nM to sustain methanogenesis, and at a maximum of 0.575 mM for phosphite oxidation to proceed (Fig. 4). This determination is consistent with DPO inhibition in our cultures when excess H_2_ was added to the headspace of the culture tubes (Figs. 4 & 5).

## Discussion

### Candidatus Phosphitivorax anaerolimi Phox-21 is an obligate lithosyntrophic phosphite oxidizer

We propose that *Candidatus* Phosphitivorax anaerolimi Phox-21’s energy metabolism and physiological dependence on a hydrogenotrophic methanogen represent a novel mode of obligate syntrophic interaction, which we have designated as “lithosyntrophy,” where the electron donor that feeds a methanogenic syntrophic interaction is inorganic, rather than organic. This expands the conceptual framework of syntrophy beyond organotrophic interactions and suggests a broader spectrum of syntrophy in anoxic ecosystems.

Several lines of evidence support the conclusion that Phox-21 shares a lithosyntrophic relationship with *Methanoculleus* sp. Phox-21’s resistance to isolation with traditional isolation methods or with substrates that have been successful for isolating facultative organosyntrophs (Supp. Fig. 1), metabolic specialization, phylogenetic relatedness to fastidious organosyntrophs (Supp. Figs. 8 and 9), indicated that Phox-21 was physiologically distinct from previously isolated DPOM FiPS-3 and DYL19 (Beaty & McInerney, 1987; Boll et al., 2016; Eichler & Schink, 1985; Ewens et al., 2021; Figueroa et al., 2017). The long-term co-enrichment of the methanogen *Methanoculleus* sp. (Fig. 1), the indirect inhibition of growth and phosphite oxidation with the specific methanogenic inhibitor BES (Fig. 2), the direct thermodynamic inhibition of hydrogenogenesis with elevated H_2_ concentrations, and the linear correlation of phosphite oxidation and methanogenesis or sulfate reduction (Fig. 3) demonstrated that Phox-21 requires a syntrophic partner that can maintain low H_2_ concentrations. Notably, the specific co-enrichment of *Methanoculleus* sp. over other hydrogenotrophic methanogens was also suggestive of an obligate syntrophic relationship: members of the genus *Methanoculleus* have a high affinity for and may be adapted to low fluxes of H_2_ produced through syntrophic oxidation, and often associate with obligately syntrophic acetate oxidizers and other species of Phosphitivorax (Barret et al., 2012; Cheng et al., 2025; Chong et al., 2002; Hao et al., 2020; Manzoor et al., 2016; Sakai et al., 2007; Schnürer et al., 1994, 1999).

These observations are consistent with the physiological differences between Phox-21 and FiPS-3 and DYL19, and aspects of our proposed model are corroborated by the metabolic model proposed by Mao and colleagues (Mao, Müller, et al., 2023, 2023). Notably, the initial use of cysteine as a reducing agent inadvertently enabled the enrichment of a previously undescribed mode of DPO that requires exogenous acetate. We initially assumed that, like FiPS-3 and DYL19, Phox-21 was a lithoautotroph coupling phosphite oxidation to carbon fixation, which was supported by the presence of the reductive glycine pathway for carbon fixation in Phox-21’s genome and necessity of CO_2_ in the growth culture (Ewens et al., 2021; Figueroa et al., 2017). However, our current study demonstrates that growth and phosphite oxidation required acetate (Supp. Figs 2 and 3), which was likely initially derived from cysteine degradation. This acetate dependency was explained by the lack of phosphoacetyltransferase (PTA) and acetate kinase (ACK) in Phox-21’s genome. In the reductive glycine pathway, these enzymes are necessary to convert glycine to acetylphosphate and then to acetate, while generating ATP via substrate-level phosphorylation (Sánchez-Andrea et al., 2020). Instead, Phox-21 converts glycine to serine and ultimately to pyruvate (Sánchez-Andrea et al., 2020). Consequently, while Phox-21’s reductive glycine pathway accounts for carbon assimilation, it cannot account for energy conservation or endogenous acetate generation and thus, we could not confidently assign CO_2_ as the terminal electron acceptor for phosphite in a single catabolic reaction.

We propose that H^+^, not CO_2_, is the electron acceptor for phosphite and that acetate is required as a co-substrate for phosphite oxidation to regenerate AMP and balance the ATP cycle via acetyl-coA synthetase (Fig. 6). This model is supported by the detection of acetyl-coA synthetase and adenylate kinase in high abundance in the proteomes of FiPS-3 and DYL19 (Mao, Fleming, et al., 2023). In FiPS-3 and DYL19, NADH generated by ptdF serves as the reducing power for lithoautotrophic CO_2_ fixation in the Wood-Ljungdahl pathway. The acetate produced by this pathway supports both carbon assimilation and ATP cycling by replenishing AMP required by PtdF. Because the reducing equivalents generated during DPO are consumed by CO_2_ fixation, these organisms may not require a syntrophic partner. In contrast, Phox-21 assimilates carbon primarily from acetate and must dispose of excess reducing equivalents through interspecies H_2_ transfer. Thus, Phox-21’s dependency on a syntrophic partner may reflect both its inability to produce acetate through an NADH-consuming process and its need to offload H_2_ generated during DPO.

Surprisingly, ptxD, the NAD^+^-dependent phosphite dehydrogenase assumed to be involved in both assimilatory phosphite oxidation (APO) and DPO (Ewens et al., 2021; Figueroa et al., 2017; Poehlein et al., 2013; Simeonova et al., 2010), was not detected in the most abundant peptides detected in our metaproteomic analysis. In concurrence with the results from a previous study on DYL19 and FiPS-3, ptdF/apdA, the NAD^+^/AMP-dependent phosphite dehydrogenase appears to be the primary enzyme involved in DPO (Mao, Fleming, et al., 2023), and indeed, PtdF was the most abundant peptide detected. In our proteomics results, PtdF and PtdC were the only enzymes encoded by the ptx-ptd gene cluster in the most abundant peptides produced by Phox-21 (Supp. Table 4). This raises the question of why evolution has conserved genes encoding two NAD^+^-dependent phosphite dehydrogenases in the same gene cluster in all identified DPOM. Perhaps PtxD is mainly used for assimilatory phosphite oxidation and can be expressed by DPOM when phosphate is extremely limiting or it may be required to maintain a NAD^+^/NADH redox balance to support anabolic reactions.

### Ecological and biogeochemical implications of lithosyntrophic DPO

Organosyntrophic interactions facilitated by interspecies H_2_ transfer are key to the methanogenic degradation of complex organic matter (McInerney et al., 2009; Schink, 1997, 2002; Stams & Plugge, 2009). Syntrophic bacteria couple the oxidation of fermentation intermediates such as organic acids and alcohols to hydrogenogenesis through proton reduction. Even under optimal growth conditions, syntrophic catabolic reactions yield very little energy, and are so thermodynamically constrained that the accumulation of small amounts of H_2_ can shift the energetics of the reaction from exergonic (Δ*G*_*r*_ < 0) to endergonic (Δ*G*_*r*_ > 0) (Jackson & McInerney, 2002; Schink, 1997). These thermodynamic constraints are overcome by the consumption of H_2_ by a syntrophic partner, which is often a methanogenic archaeon. Interspecies H_2_ transfer allows for the complete degradation of fermentation intermediates that neither syntrophic partner can degrade alone.

In lithosyntrophy, phosphite oxidation drives hydrogenotrophic methanogenesis (Fig. 7A). The discovery of lithosyntrophic phosphite oxidation reveals a previously unrecognized mechanism by which inorganic compounds contribute to methanogenesis and carbon cycling in anoxic environments. Microbial methane production is canonically assumed to be driven by complex organic matter degradation to H_2_/CO_2_, formate, acetate, and methylated compounds (McInerney et al., 2009; Schink, 2002; Thauer et al., 2008) or by geologically produced H_2_ in subsurface environments (Boyd et al., 2024; Lollar et al., 2014; Stevens & McKinley, 1995). However, the discovery of lithosyntrophic phosphite oxidation demonstrates that inorganic compounds can serve as an energy source in obligate syntrophic interactions, through the coupling of phosphite oxidation to hydrogenotrophic methanogenesis (Table 1, Rxns. 4 and 17). This expands the known metabolic diversity of syntrophic interactions and establishes an unrecognized link between the phosphorus and carbon cycles. Archaeal methanogenesis in anoxic environments is estimated to produce more than half of annual methane emissions (Schwietzke et al., 2016), and lithosyntrophy may serve as a major but previously unaccounted driver of methanogenesis.

**Figure 7.**
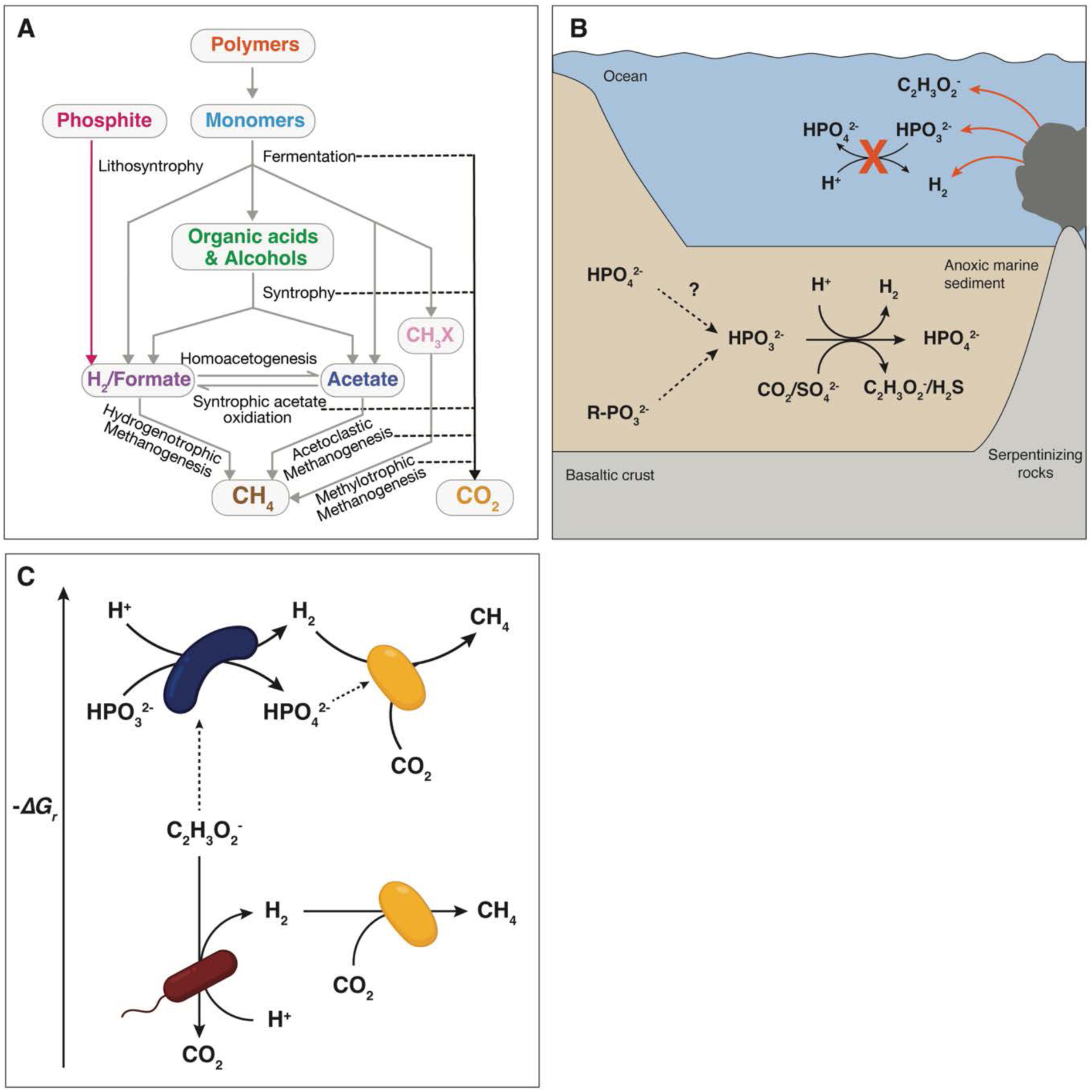
Geochemical and ecological contexts of lithosyntrophic phosphite oxidation. A) Electron flow in organotrophic and lithosyntrophic carbon degradation. While organosyntrophy drives methanogenesis through hydrogen production from organic acids or alcohols, lithosyntrophy similarly supports methanogenesis via hydrogen production from phosphite oxidation. B) Mechanisms of phosphite production and geochemical conditions that may support or inhibit lithosyntrophic DPO. Serpentinization produces hydrogen, phosphite, acetate, and highly reducing conditions, but high hydrogen concentrations are inhibitory to lithosyntrophic DPO. In anoxic marine sediments, phosphite may be produced through phosphate reduction or phosphonate degradation and can be oxidized through lithosyntrophic DPO, acetogenic DPO, or sulfidigenic DPO. Red lines indicate abiotic processes, dotted lines indicate hypothetical processes, and solid black lines indicate known microbial metabolic reactions. C) Proposed ecological and energetic advantages for hydrogenotrophic methanogens (yellow) that partner with lithosyntrophic DPOM (blue). Compared to acetoclastic methanogens or syntrophy with acetate oxidizers (red), hydrogenotrophic methanogens in lithosyntrophic associations gain more energy per mole of carbon or per mole of electrons transferred, and additionally acquire bioavailable phosphorus. Solid black lines indicate dissimilatory reactions and dashed lines indicate assimilatory processes.

A key outstanding question is where and how phosphite is produced in present day environments, which limits our understanding of the environmental distribution and ecological relevance of lithosyntrophic DPOM. FiPS-3, DYL19, and Phox-21 were isolated or enriched from engineered environments (wastewater treatment sludge) or impacted environments (Canale Grande, Venice, Italy). In wastewater treatment sludge, phosphite concentrations have been measured at 0.157 – 0.278 µM (Sadeghi & Jackson, 2024; Yu et al., 2015). Few measurements of phosphite in natural environments have been reported (Han et al., 2013; Pech et al., 2009), but in freshwater samples from Florida (rivers, ponds, and swamps), reduced oxidation state phosphorus (phosphite and hypophosphite) accounted for 10-20% of the total phosphorus pool (M. A. Pasek et al., 2014). Higher phosphite concentrations were detected in the stagnant waters (ponds/swamps) compared to the river samples. The pond environments in this study are impacted by anthropogenic activity, while the swamp represents a natural environment. Like wastewater treatment sludge, the stagnant waters where phosphite was detected by Pasek et al. are likely anoxic and nutrient rich. The source of phosphite in these systems is unclear, but it may be produced through phosphate reduction or phosphonate degradation in the course of organic matter catabolism in high carbon, reducing environments (Fig. 7B).

In marine and terrestrial serpentinite muds, phosphite was detected at 23-60% of total P (50-100 µM) (M. A. Pasek et al., 2022). The process of serpentinization generates H_2_ which can react with CO_2_ to produce fluids enriched in small organics like acetate (Preiner et al., 2018, 2020; Schwander et al., 2023). ptdF was detected in metagenomic datasets from serpentinizing sites at the Lost City Hydrothermal Field, suggesting that microorganisms with the potential for DPO exist at these sites (Boden et al., 2025). While the presence of acetate, phosphite, and the key marker gene for DPO are promising, serpentinizing fluids are characterized by high H_2_ concentrations, with concentrations measured at up to 15 mM at the Lost City Hydrothermal Field (Kelley et al., 2005; Proskurowski et al., 2006). While it is possible that acetogenic DPO could serve as a microbial metabolism at serpentinizing sites, high background H_2_ concentrations in vent fluids would likely be inhibitory to lithosyntrophic DPOM (Fig. 7B).

In nutrient-limited environments, such as deep subsurface sediments, lithosyntrophic phosphite oxidizers may play important ecological roles in transferring both reducing equivalents (in the form of H_2_) and bioavailable phosphate to their surrounding microbial communities (Fig. 7C). Because phosphate has been a limiting nutrient for primary production across geologic time (Benitez-Nelson, 2000; Reinhard et al., 2017), the ability to perform DPO, particularly in environments where phosphite is the dominant phosphorus species, may give lithosyntrophic DPOM a competitive advantage, potentially acting as keystone species that shape microbial ecosystem structure and function. Because acetate is required as a co-substrate for DPO, lithosyntrophic DPOM likely compete for acetate with acetoclastic methanogens and syntrophic acetate oxidizers.

Under simulated marine sediment conditions, at 4 °C, with phosphite and acetate concentrations set at 1 µM, lithosyntrophic DPO and syntrophic acetate oxidation yield virtually the same amount of energy per mole of H_2_ (Supp. Fig. 10). When normalized per mole of carbon with the concentration of H_2(aq)_ set at 10 nM, however, methanogenic DPO by lithosyntrophic DPO in association with hydrogenotrophic methanogens (Table 1, Rxn. 17) yields −16.3 kJ/mol carbon (−2.04 kJ/mol e^−^), while syntrophic acetate oxidation (Table 1, Rxn. 18) coupled to hydrogenotrophic methanogenesis (the net reaction of which can be represented as acetoclastic methanogenesis; Table 1, Rxn. 19) yields only −1.1 kJ/mol carbon (−0.53 kJ/mol e^−^). Under these conditions, methanogenic DPOM partnered with hydrogenotrophic methanogens should outcompete syntrophic acetate oxidizers partnered with hydrogenotrophic methanogens and acetoclastic methanogens on an energetic basis (Fig. 7C). It might be advantageous for a hydrogenotrophic methanogen to form a lithosyntrophic partnership with a DPOM because it would receive a supply of electron donor and bioavailable phosphate.

It has been proposed that phosphite was more abundant early in Earth’s history because of highly reducing conditions and could have been produced abiotically through potential input from meteoric phosphite minerals such as schreibersite, (Fe,Ni)_3_P, or phosphate reduction by lightning, ferrous iron, or H_2_ during serpentinization or high-grade metamorphism in the Archaean (Baidya et al., 2024; Herschy et al., 2018; M. A. Pasek, 2008; M. A. Pasek et al., 2013, 2022; M. Pasek & Block, 2009). Lithosyntrophic phosphite oxidation coupled to hydrogenotrophic methanogenesis (Table 1,. 17) provides a model for how primordial microbial communities may have been sustained exclusively by geologically-sourced substrates in electron acceptor-limited, anoxic ecosystems (Nealson et al., 2005). These two metabolisms require substrates that can be produced geologically (phosphite, protons, CO_2_, and acetate) which could have been available nutrients early in Earth’s history. In our proposed metabolic model for lithosyntrophic DPO, H_2_ is produced through electron confurcation, and energy is conserved through substrate-level phosphorylation and pyrophosphate hydrolysis coupled to ion translocation (Fig. 6), which is suggestive of evolutionarily ancient metabolisms (Baltscheffsky et al., 1998; Baykov et al., 2013; Ferry, 2006; Martin, 2020; Martin & Thauer, 2017; Müller et al., 2018; Nicholls et al., 2023).

Phylogenomic reconstructions of the ptx-ptd gene cluster estimate that DPO emerged ∼3.2 Ga in the most recent common ancestor of the Firmicutes and Desulfobacterota (Ewens et al., 2021). While molecular clock analyses by Boden et al. 2024 estimate the earliest evolution of DPO in the Firmicutes around 2.3 Ga, these predictions were based solely on the presence of ptxD, which does not appear to be involved in DPO (Boden et al., 2024). Instead, ptdF and ptdC, which were detected in high abundance in the proteomes of Phox-21, FiPS-3, and DYL19, should be considered as the key marker genes of DPO, while ptxD is likely primarily indicative of assimilatory phosphite oxidation (Mao, Fleming, et al., 2023). In either case, the origin of DPO likely occurred after the evolution of hydrogenotrophic methanogenesis (∼4.1 Ga - 3.7 Ga) and predated the emergence of acetoclastic methanogenesis (∼250 Ma) (Battistuzzi et al., 2004; Rothman et al., 2014). During the Archaean, when phosphite was likely more abundant, lithosyntrophic DPO would have been a sink for acetate. Following the Great Oxidation Event (GOE, 2.50 – 2.20 Ga), the progressive oxygenation of the environment would have abiotically diminished phosphite availability, reducing the ecological prevalence of DPO, and, in turn, decreasing competition for acetate. In this context, the later emergence of acetoclastic methanogenesis may reflect adaptation to more oxidizing conditions in which acetate became a more accessible methanogenic substrate.

## Conclusion

These findings broaden our understanding of the role of inorganic compounds in anaerobic carbon cycling by revealing previously unrecognized lithosyntrophic interactions and point to the potential existence of similar interactions involving other inorganic substrates. The demonstration that phosphite oxidation can drive hydrogenotrophic methanogenesis redefines the role of phosphorus in anoxic ecosystems and establishes a new metabolic and biogeochemical link between the phosphorus and carbon cycles. Lithosyntrophic DPOM may act as keystone species in certain environments, serving as producers of reducing equivalents and sources of bioavailable phosphorus. Together, these results suggest that lithosyntrophic DPO may play a critical role in supporting primary production, driving methanogenesis, and redistributing bioavailable phosphorus in anoxic, high-carbon environments, engineered systems, electron acceptor-limited environments, the modern deep biosphere, and in potentially in early Earth environments.

## Methods

### Thermodynamic calculations

The Gibbs energy of reaction *ΔG*_*r*_, was calculated for 19 reactions involved in the DPO enrichment culture (Table 1, Supp. Table 1), using

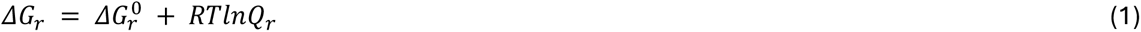

where 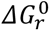 is the standard state Gibbs energy, *R* refers to the universal gas constant, *T* represents the temperature in Kelvin, and *Q_r_* denotes the reaction quotient. Values of *ΔG*^0^ were calculated for each reaction at 37 °C and 1 bar using the revised Helgeson-Kirkham-Flowers (HKF) equations of state (Helgeson et al., 1981; Shock & Helgeson, 1988; Tanger & Helgeson, 1988) using the “subcrt” command from the R software package CHNOSZ v2.1.0 (Dick, 2019). Thermodynamic data in CHNOSZ are sourced from the OrganoBioGeoTherm database, which are derived from a number of sources documented in https://chnosz.net/download/refs.html. Values represented by the activity quotient, *Q_r,_* were calculated with

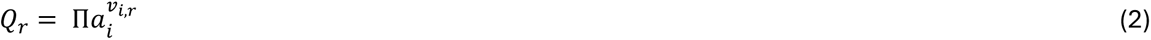

where *a*_*i*_ corresponds to the activity of the *i*th species raised to its stoichiometric coefficient, *v*_*i*,*r*_, in the *r*th reaction, which is positive for products and negative for reactants. Activity quotients were calculated from the chemical composition of the cultivation medium (Supp. Table 2, described below) with a range of H_2_(aq) concentrations using the “ae.speciate” command from the aqueous speciation package AqEquil v0.18.0 (Supp. Tables 3 and 4) (Boyer et al., 2024). Intracellular concentrations of ATP (2.2 mM), ADP (0.83 mM), and AMP (0.3 mM) were sourced from Thauer et al., 1977. Intracellular concentrations of reduced and oxidized ferredoxin, NAD^+^, and NADH were assumed to be 100 µM. The standard state Gibbs energy for ferredoxin oxidation was calculated from its redox potential (Buckel & Thauer, 2013). Henry’s Law constants (mol L^−1^ atm^−1^) for hydrogen and methane were calculated at 37 °C using the equilibrium constants of the gas dissolution reaction for each compound (Table 1, Rxns. 2 and 3). To model the energetics of these reactions under simulated marine sediment conditions, aqueous speciation and energetics calculations were carried out at 4 °C with phosphite and acetate concentrations set at 1 µM, major cations and anions set at marine concentrations, with the concentrations of other products and reactants as described above (Supp Table 3).

### Media and cultivation conditions

Highly enriched phosphite oxidizing enrichment cultures were originally inoculated and maintained by Ewens et al., 2021. Briefly, anoxic, HEPES buffered basal medium (Supp. Table 2) was prepared and dispensed under N_2_/CO_2_ (80:20). All chemicals were sourced from Sigma-Aldrich. Medium amendments were added to the basal medium from sterile, anoxic 1 M stocks. The medium was reduced with 3 mM L-cysteine HCl and 10 mM sodium phosphite dibasic pentahydrate was added as the electron donor and sole phosphorus source. Enrichment cultures were spiked with an additional 10 mM sodium phosphite after two weeks of growth at 37 °C and passaged with a 10% inoculation volume after one month. Growth was measured as optical density at 600 nm (OD_600_) with a Genesys 20 Visible spectrophotometer (Thermo Scientific).

L-cysteine HCl was replaced with 3 mM sodium acetate and 1 mM sodium sulfide nonahydrate. For isolation experiments, phosphite was replaced with 10 mM sodium phosphate monobasic and the media was amended with 10 mM crotonic acid or acetoin from sterile, anoxic stock solutions. For conditions with excess H_2_, 10 mL of H_2_ gas was added to the headspace of a Balch tube containing 10 mL of culture. Tubes with excess H_2_ were incubated shaking (250 rpm) and positioned at a slant to increase the dissolution of H_2_ into the liquid medium. Methanogenesis was inhibited with 1 mM 2-bromoethanesulfonate (BES). The methanogenic DPO culture was grown under sulfate-reducing conditions (10 mM sulfate) in the presence of BES by adding the model hydrogenotrophic sulfate reducer, *Desulfovibrio desulfuricans* G11 at 50% of the inoculum.

### Isolation and characterization of Methanoculleus sp

To isolate the methanogen, *Methanoculleus* sp., the anoxic basal medium was amended with 10 mM sodium phosphate monobasic and H_2_ was added at 20% of the headspace volume. Methane was measured as described below and the culture was isolated by serial dilution to extinction. Sample purity was assessed by 16S rRNA amplicon sequencing and metagenomic sequencing, as described below. To determine which methanogenic substrates could be used, *Methanoculleus* was grown at 37 °C with 20:80 H_2_/CO_2_ (shaking at a slant, 250 rpm), with 10 mM sodium formate, or with 10 mM sodium acetate.

### Sampling and measurements of ions and gases

300 µL culture was collected and filtered through a 0.2 µm nylon syringe filter for analytical measurements and stored at 4 °C until analysis. Phosphite, phosphate, and sulfate concentrations were measured via ion chromatography using a Dionex ICS 2100 with a Dionex IonPac AS 16 column (4 x 250 mm) (Thermo Fisher Scientific) as described in (Figueroa et al., 2017). Headspace methane concentrations were measured via gas chromatography using an Agilent gas chromatograph model 7890A (G344OA) with a Supelco SP-2380 Fused Silica Capillary column. Methane was measured using a flame ionization detector (FID) at 300 °C, a column temperature of 50 °C. Headspace hydrogen concentrations were measured by gas chromatography with a reducing compound photometer using a Peak Performer 1 reductive gas analyzer (Peak Laboratories) with a bed temperature of 265 °C, a column temperature of 105 °C, and N_2_ as the carrier gas.

### Stable isotope probing and metaproteomics

50 mL cultures were amended with either 10 mM ^13^C-sodium bicarbonate (Sigma Aldrich), 3 mM sodium acetate-2-^13^C, or 3 mM sodium acetate-2-^13^C without CO_2_ and bicarbonate, and incubated for one week at 37°C. ^13^C incorporation into headspace methane and CO_2_ was measured by gas chromatography-mass spectrometry (GC-MS; Agilent) with a GasPro column on day 0 and day 7 of incubation. The ratio of ^13^CH_4_ to ^12^CH_4_ on day 7 was calculated and significance testing was performed using analysis of variance (ANOVA) and Tukey’s honestly significant difference (HSD) test.

After one week of incubation, the entire culture (50 mL) was pelleted for tryptic digestion by centrifuging at 16,873 relative centrifugal force (rcf) for 10 minutes. The pellet was washed and resuspended in 100 mM NH_4_HCO_3_, pH 7.5, and transferred to a microcentrifuge tube. The pellet was centrifuged again and resuspended in 50 µL of 100 mM NH_4_HCO_3_. 25 µL of 0.2% RapiGest SF surfactant (Waters 186001861) was added to make the proteins more accessible for alkylation and digestion and the samples were heated for 15 minutes at 80 °C. 2.5 µL of 100 mM dithiothreitol (15.4 µg/µL) was added and the samples were heated for 30 minutes at 60 °C. The samples were cooled to room temperature and 7.5 µL of 100 mM iodoacetamide (55.5 µg/µL) was added to alkylate cysteines. The tubes were incubated at room temperature in the dark for 30 minutes. 5 µL of Promega trypsin (1 µg/µL) was added and the samples were incubated overnight at 37 °C. Following the trypsin digestion, 10 µL of 5% trifluoroacetic acid was added to hydrolyze the RapiGest. The samples were incubated for 90 minutes at 37 °C, then centrifuged for 30 minutes at 14,000 RPM at 4 °C. 30 µL of the supernatant was transferred to a sample vial.

Triplicate trypsin-digested protein samples of unlabeled controls, ^13^C-bicarbonate, and ^13^C-acetate were analyzed with an Acquity M-class ultra-performance liquid chromatography (UPLC) system that was connected in line with Synapt G2-Si mass spectrometer (Waters, Milford, MA) as described previously (Grant et al., 2022). Data were acquired and analyzed using MassLynx (v4.1, Waters) and Progenesis QI for Proteomics software (v4.2, Waters Nonlinear Dynamics). Data were searched against the translated protein sequence of the metagenome (described below).

### DNA extraction, amplicon and metagenomic sequencing, and bioinformatics

1.5 mL of culture was collected for DNA extraction and centrifuged for 10 minutes at 16,873 rcf. Supernatant was removed and pellets were stored at −20 °C until DNA extraction. DNA was extracted for 16S rRNA community sequencing using enzymatic digestion followed by ethanol precipitation. Pellets were resuspended in 200 µL enzymatic lysis buffer (20 mM Tris-HCl pH 8.0, 2 mM sodium EDTA, 1.2% Triton X-100) and transferred to a 96 well plate. 20 µL Proteinase K (20 mg/mL) and 200 µL of 4M guanidine HCl was added to each well and plates were incubated overnight at 55 °C. 4 µL of RNAse A (100 mg/mL) was added to each well and plates were incubated for 2 hours at room temperature. 200 µL of 100% ethanol was added and the contents of each well were transferred to a 96-well plate filter. Columns were washed with 400 µL of 70% ethanol and DNA was eluted with 100 µL of TE buffer (100 mM Tris, 10 mM EDTA, pH 8.0). The V4/V5 16S rRNA gene region was amplified via PCR using the 515F/926R primers (Parada et al., 2016), but with in-line dual Illumina indexes (Price et al., 2018; Sharpless et al., 2022). Amplicons were sequenced on an Illumina MiSeq (Illumina, San Diego, CA, USA) with 2×300 bp Illumina v3 reagents. Custom Perl scripts implementing PEAR (Zhang et al., 2014) were used read merging, USearch (Edgar, 2010) for filtering reads with more than one expected error and demultiplexing using inline indexes, and UNoise (Edgar, 2016) for filtering rare reads and chimeras. 16S sequences in the relative abundance table were searched against the NCBI RefSeq database and matched metagenome-assembled genomes (MAGs, described below) to assign taxonomy.

DNA was extracted for metagenomic sequencing using the Qiagen DNeasy UltraClean DNA extraction kit following manufacturer’s protocols. Metagenomic libraries were prepared and sequenced on a NovaSeqX platform by Novogene Corporation Inc. Metagenomic raw reads were trimmed using Sickle (Joshi & Fass, 2011/2011) to remove low quality reads (q<20). Triplicate metagenomes (biological replicates) were co-assembled using MEGAHIT (Li et al., 2015) and contigs shorter than 1000 base pairs were eliminated using the “anvi-script-reformat-fasta” command from the software package anvi’o (Eren et al., 2021). Raw reads were mapped to the assembly using bowtie2 and sorted and indexed using samtools (Danecek et al., 2021; Langmead & Salzberg, 2012). Genome binning of assembled reads was performed using MetaBat, MetaBAT 2 (Kang et al., 2019), concoct (Alneberg et al., 2014), and MaxBin 2.0 (Wu et al., 2016). An optimized, non-redundant set of MAGs was selected using DAS Tool (C. M. K. Sieber et al., 2018). MAG quality was assessed using CheckM2 (Chklovski et al., 2022) and taxonomic identity was determined using GTDBTK v2.4.1, release 226 (Chaumeil et al., 2022). MAG genome sequences were translated to protein sequences using Prokka (v1.14.6) (Seemann, 2014), and protein sequences for each MAG were concatenated into a single file to use for peptide identification from the metaproteome.

A phylogenomic tree of the phylum Desulfobacterota was constructed using Phox-21 MAGs from this study and from (Ewens et al., 2021; Figueroa et al., 2017) using the GTDBTK de novo workflow (Chaumeil et al., 2022). A 16S rRNA phylogenetic tree of Phox-21, *Desulfotignum phosphitoxidans* FiPS-3, *Phosphitispora fastidiosa* DYL19, and Phox-21’s closest relatives was constructed using the top 1000 sequence matches from the NCBI core nucleotide (core_nt) database and the top 10 sequence matches from the NCBI 16S rRNA type strain database. The 16S rRNA sequence for *Sulfurovum aggregans* was used as an outgroup. Sequences were dereplicated using the rmdup command from SeqKit (Shen et al., 2016), aligned using MUSCLE (Edgar, 2004), trimmed using trimAl (Capella-Gutiérrez et al., 2009), and a tree was constructed using IQ-Tree (Nguyen et al., 2015). Trees were visualized using iTOL (Letunic & Bork, 2021).

## Author contributions

H.S.A., M.E.W, H.K.C., and J.D.C. designed research; H.S.A., M.E.W, R.H., V.C.L., and J.D.R. performed the research; A.T.I. performed mass spectrometry analyses and assisted with proteomic data analysis; H.S.A. analyzed the data; H.S.A. and J.D.C. wrote the paper. All authors reviewed and approved the final version of the manuscript. The authors declare no conflicts of interest.

## Data availability

Metagenomic reads for the enrichment culture containing cysteine are available in the NCBI BioProject accession no. PRJNA655520. Metagenomic reads and 16S rRNA short read sequences for the enrichment culture containing acetate and sulfide are available in the NCBI BioProject accession no. PRJNA1305015.

## Acknowledgements

This work was supported by Energy & Biosciences Institute-Shell (EBI-Shell) research funding to JDC. H.S.A. was supported by the EBI-Shell Postdoctoral Fellowship. A mass spectrometer was purchased with support from the National Institutes of Health (grant number 1S10OD020062-01). We thank Yi Liu for laboratory support and Grayson Boyer for assistance with aqueous speciation calculations.

---

| Number | Reaction | Description |
| --- | --- | --- |
| 1 | $4\text{HPO}_3^{2-} + 2\text{HCO}_3^- + \text{H}^+ \rightarrow 4\text{HPO}_4^{2-} + \text{C}_2\text{H}_3\text{O}_2^-$ | Acetogenic DPO |
| 2 | $\text{CH}_{4(g)} \leftrightarrow \text{CH}_{4(aq)}$ | Methane dissolution |
| 3 | $\text{H}_{2(g)} \leftrightarrow \text{H}_{2(aq)}$ | Hydrogen dissolution |
| 4 | $4\text{HPO}_3^{2-} + \text{HCO}_3^- + \text{H}_2\text{O} + \text{H}^+ \rightarrow 4\text{HPO}_4^{2-} + \text{CH}_{4(aq)}$ | Lithosyntrophic DPO coupled to hydrogenotrophic methanogenesis |
| 5 | $\text{HPO}_3^{2-} + \text{H}_2\text{O} \rightarrow \text{HPO}_4^{2-} + \text{H}_{2(aq)}$ | Lithosyntrophic DPO |
| 6 | $4\text{HPO}_3^{2-} + \text{SO}_4^{2-} + \text{H}^+ \rightarrow 4\text{HPO}_4^{2-} + \text{HS}^-$ | Lithosyntrophic DPO coupled to sulfate reduction |
| 7 | $4\text{H}_{2(aq)} + \text{HCO}_3^{2-} + \text{H}^+ \rightarrow \text{CH}_{4(aq)} + 3\text{H}_2\text{O}$ | Hydrogenotrophic methanogenesis |
| 8 | $4\text{H}_{2(aq)} + \text{SO}_4^{2-} + \text{H}^+ \rightarrow \text{HS}^- + 4\text{H}_2\text{O}$ | Hydrogenotrophic sulfate reduction |
| 9 | $\text{HPO}_3^{2-} + \text{AMP} + \text{H}^+ \rightarrow \text{H}_2 + \text{ADP}$ | Lithosyntrophic DPO, low energy electrons (ptdF + Hnd + Rnf) |
| 10 | $2\text{HPO}_3^{2-} + 2\text{NAD}^+ + 2\text{AMP} \rightarrow 2\text{NADH} + 2\text{ADP}$ | ptdF |
| 11 | $\text{NADH} + 2\text{Fd}_{red} + 3\text{H}^+ \rightarrow \text{NAD}^+ + 2\text{Fd}_{ox} + 2\text{H}_{2(aq)}$ | HndABCD |
| 12 | $\text{NADH} + 2\text{Fd}_{ox} \rightarrow \text{NAD}^+ + 2\text{Fd}_{red} + \text{H}^+$ | Rnf |
| 13 | $2\text{ADP} \rightarrow \text{ATP} + \text{AMP}$ | Adenylate kinase |
| 14 | $\text{Acetate} + \text{ATP} + \text{CoA} + \text{H}_2\text{O} \rightarrow \text{AMP} + \text{P}_2\text{O}_7^{4-} + \text{Acetyl-CoA} + 2\text{H}^+$ | Acs |
| 15 | $\text{P}_2\text{O}_7^{4-} + \text{H}_2\text{O} \rightarrow 2\text{HPO}_4^{2-}$ | Pyrophosphatase |
| 16 | $\text{NADH} + \text{H}^+ \rightarrow \text{NAD}^+ + \text{H}_{2(aq)}$ | NADH oxidation coupled to proton reduction |
| 17 | $4\text{HPO}_3^{2-} + \text{HCO}_3^- + 4\text{AMP} + 5\text{H}^+ \rightarrow \text{CH}_{4(aq)} + 4\text{ADP} + 3\text{H}_2\text{O}$ | Lithosyntrophic DPO coupled to methanogenesis, low energy electrons |
| 18 | $\text{C}_2\text{H}_3\text{O}_2^- + 4\text{H}_2\text{O} \rightarrow 2\text{HCO}_3^- + 4\text{H}_{2(aq)} + \text{H}^+$ | Syntrophic acetate oxidation |
| 19 | $\text{C}_2\text{H}_3\text{O}_2^- + \text{H}_2\text{O} \rightarrow \text{CH}_{4(aq)} + \text{HCO}_3^-$ | Acetoclastic methanogenesis |

