## Supplementary Figures for "Lithosyntrophy: Obligate syntrophy in a phosphite-oxidizing, methanogenic culture"

**Supplemental figures**

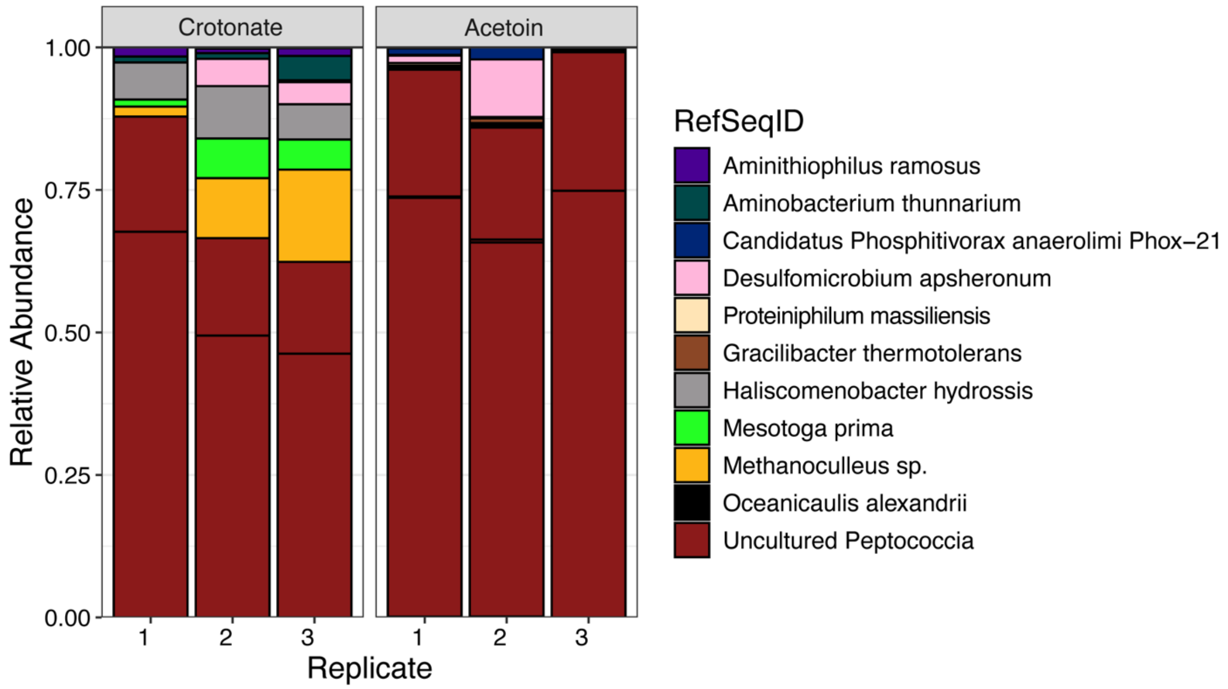


**Supplemental Figure 1:** Relative abundances of microbial taxa from enrichment cultures A) with crotonate and B) with acetoin. Each bar represents a replicate culture. Taxonomic identification is based on the V4/V5 16S rRNA gene region.

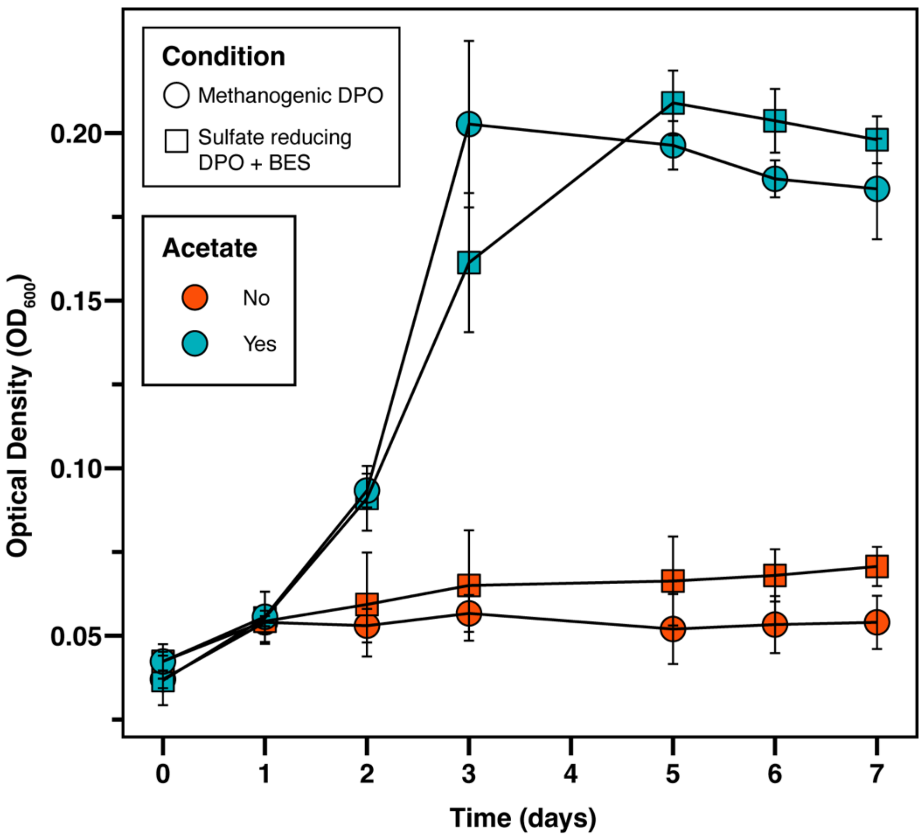


**Supplemental Figure 2:** Acetate-dependent DPO activity. Growth ($OD_{600}$) of methanogenic and sulfidigenic phosphite-oxidizing enrichment cultures with and without 3 mM acetate. Error bars represent SD of triplicate cultures.

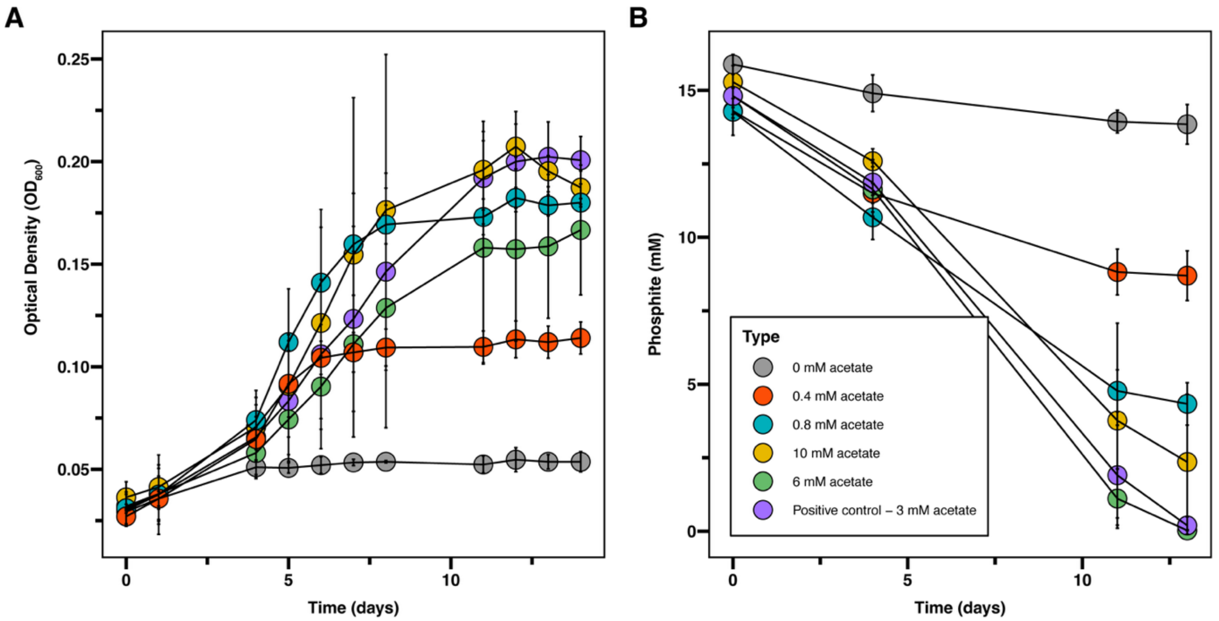


**Supplemental Figure 3:** Growth (A) and phosphite concentration (B) of methanogenic phosphite-oxidizing enrichment cultures with a gradient of acetate concentrations. Error bars represent SD of triplicate cultures.

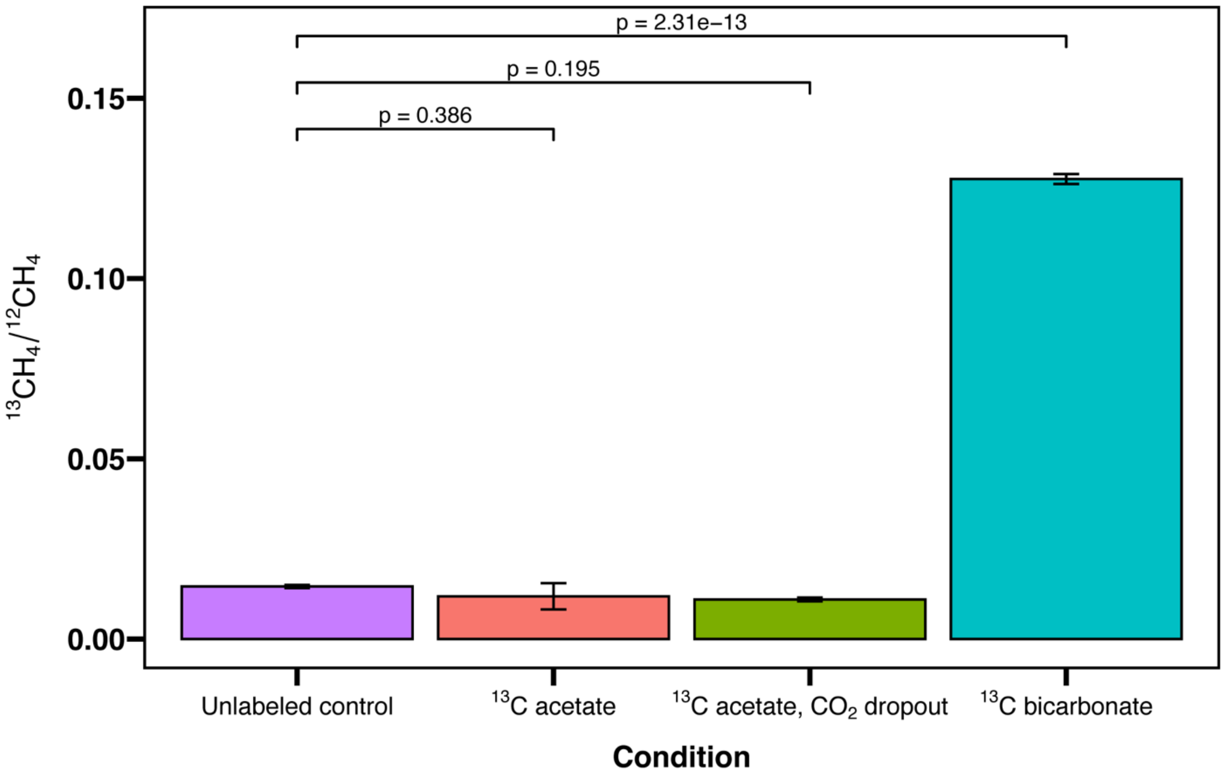


**Supplemental Figure 4:** Ratio of $^{13}C$:$^{12}C$ in methane measured in methanogenic phosphite-oxidizing enrichment cultures with different carbon sources. Statistical significance was performed using ANOVA and Tukey's HSD test.

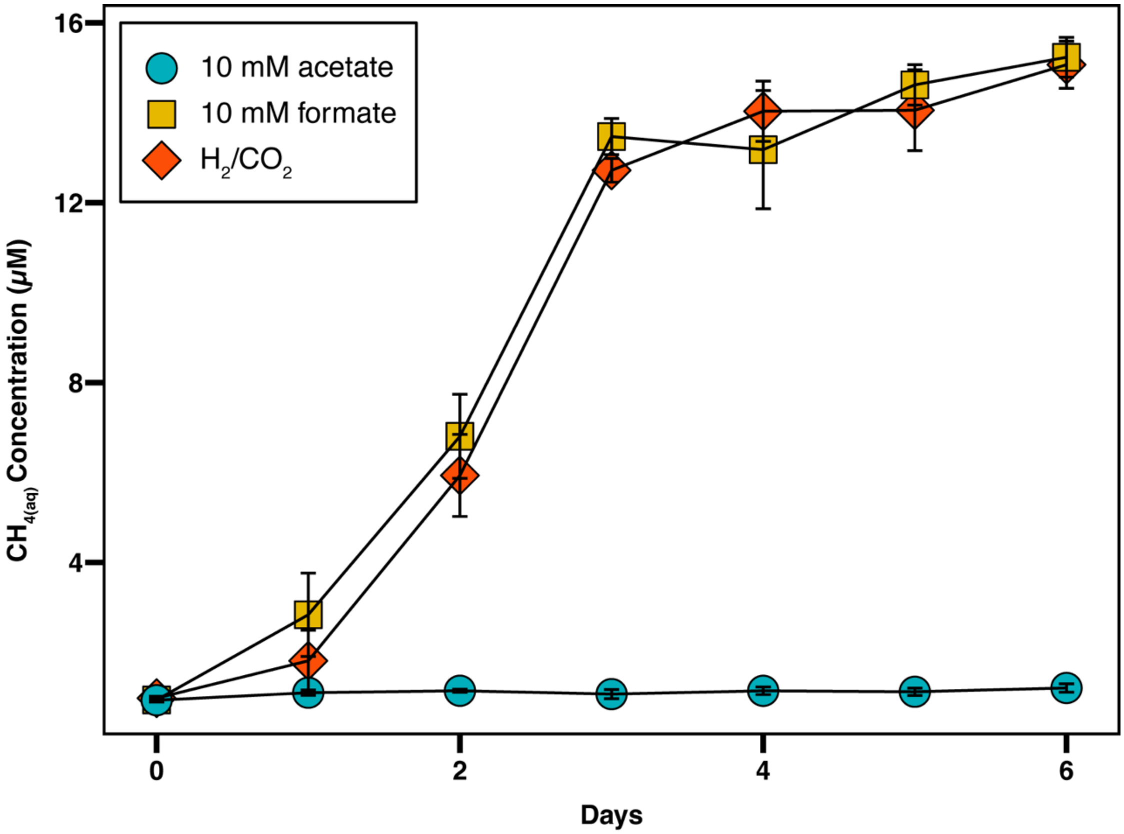


**Supplemental Figure 5:** Methane concentration (μM) in *Methanoculleus* sp. cultures grown on 10 mM sodium acetate, 10 mM sodium formate $H_2/CO_2$, (20:80). Headspace methane concentrations measured via gas chromatography were used to calculate the concentration of aqueous methane. Error bars represent SD of triplicate cultures.

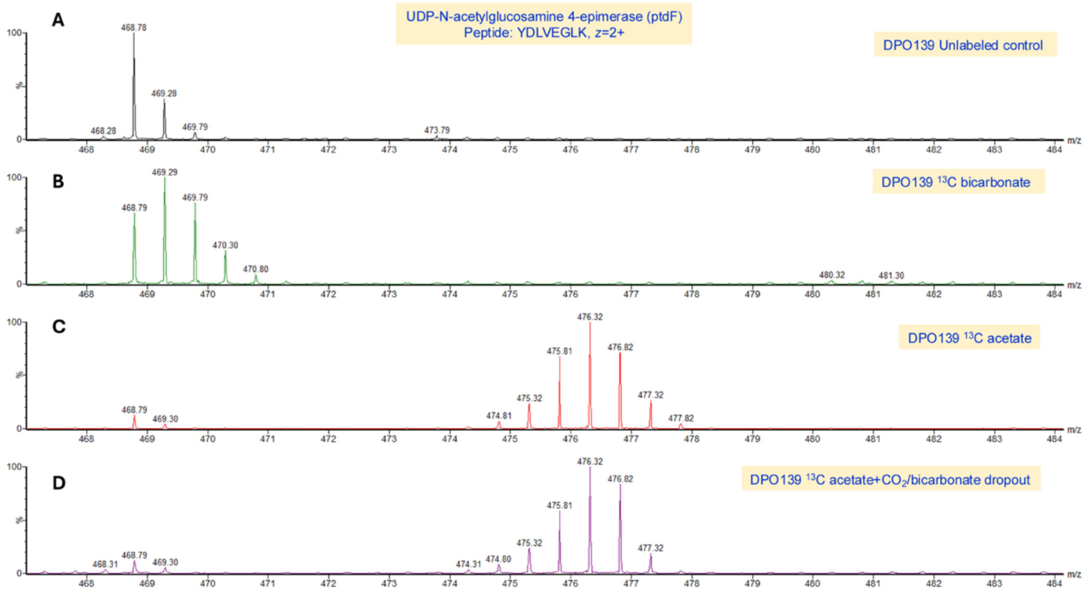


**Supplemental Figure 6:** Mass spectra of peptide YDLVEGLK from PtdF, the most abundant peptide produced by Phox-21, from A) the unlabeled control condition, B) the condition amended with $^{13}C$ bicarbonate, C) the condition amended with $^{13}C$ acetate, and D) the condition amended with $^{13}C$ acetate with $CO_2$ and bicarbonate dropped out.

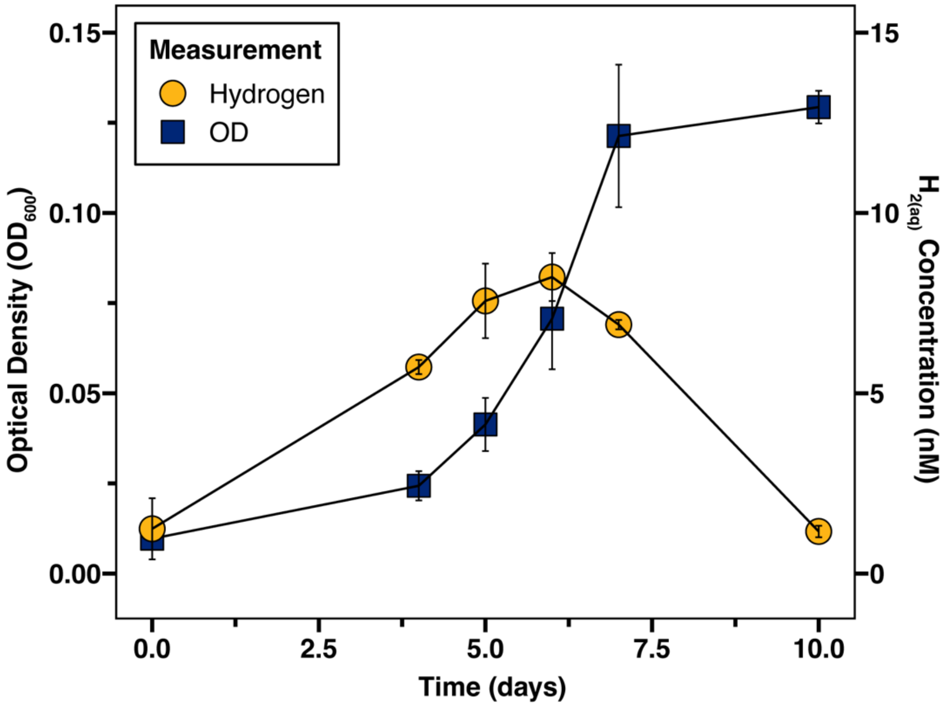


**Supplemental Figure 7:** Growth ($OD_{600}$, blue squares) and aqueous hydrogen concentration (yellow circles) of methanogenic phosphite-oxidizing enrichments. Measured headspace hydrogen concentrations were used to calculate the aqueous hydrogen concentration in the media. Error bars represent SD of triplicate cultures.

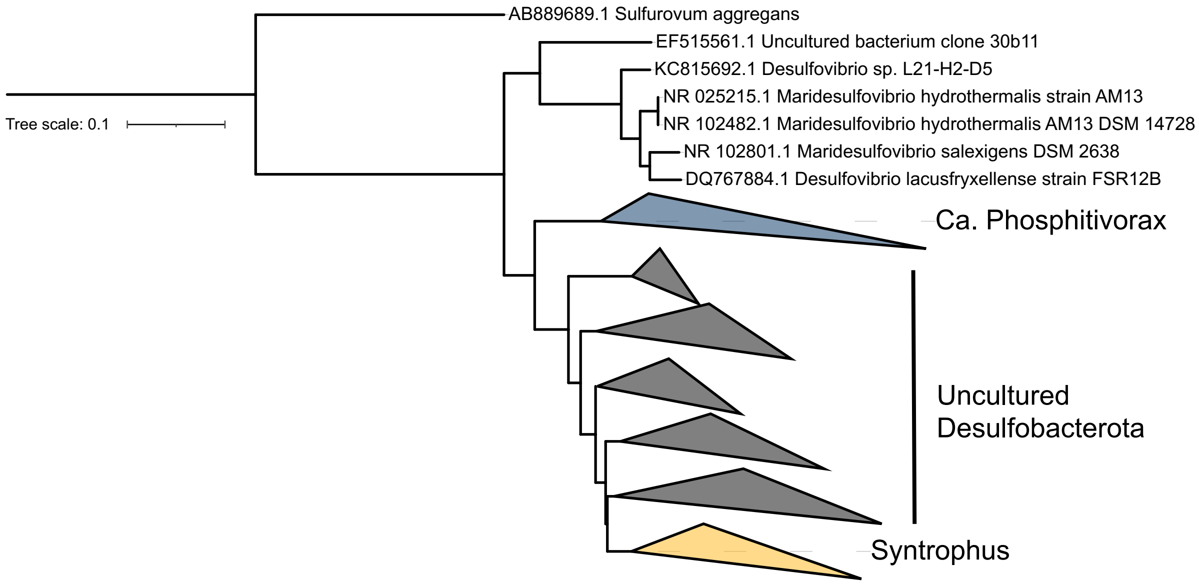


**Supplemental Figure 8:** Phylogenetic tree of 16S rRNA sequences from Phox-21 and close relatives from the NCBI core nucleotide and 16S rRNA type strain database.

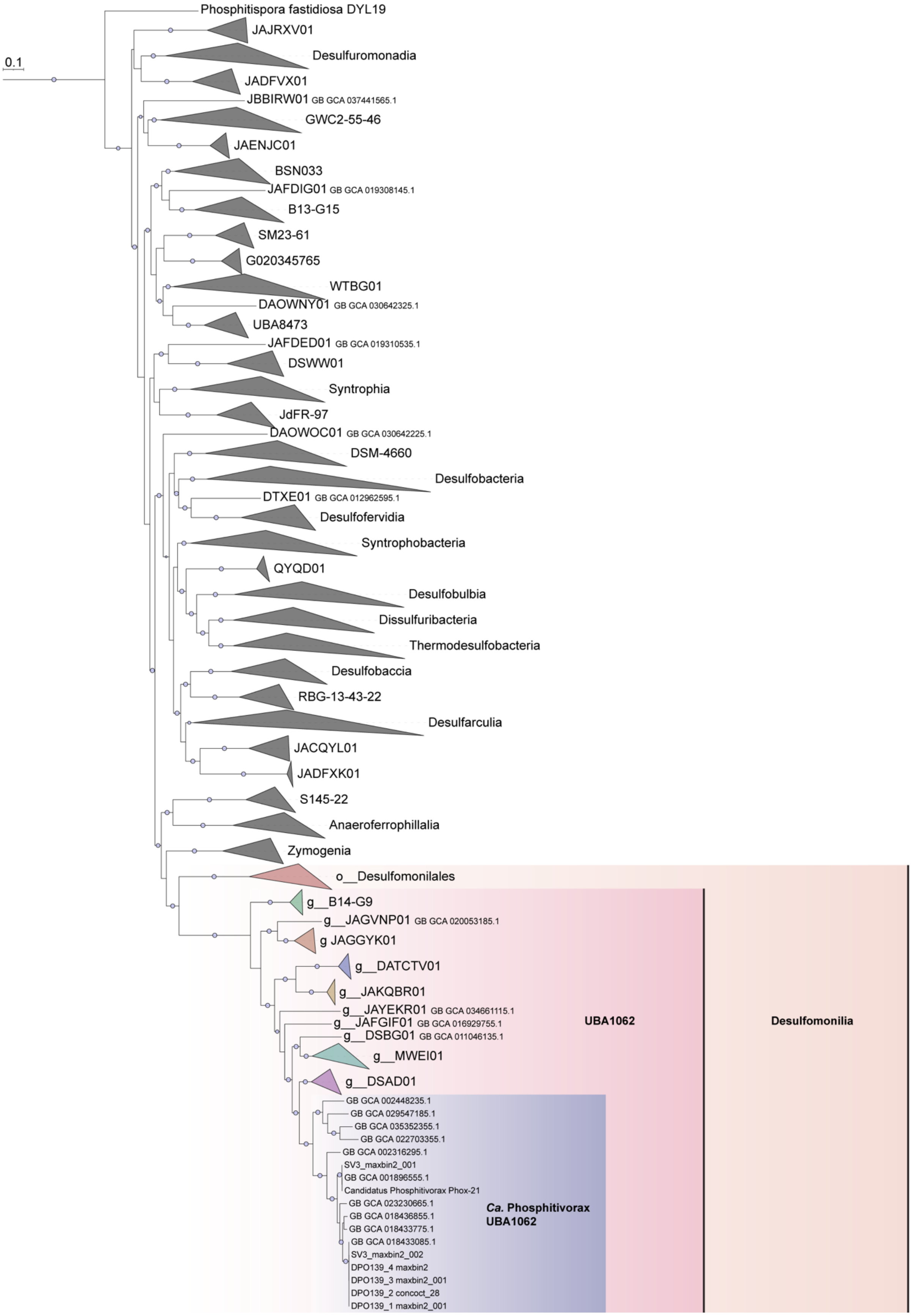
0.1
Phosphitispora fastidiosa DYL19
JAJRXV01
Desulfuromonadia
JADFVX01
JBBIRW01 GB GCA 037441565.1
GWC2-55-46
JAENJC01
BSN033
JAFDIG01 GB GCA 019308145.1
B13-G15
SM23-61
G020345765
WTBG01
DAOWNY01 GB GCA 030642325.1
UBA8473
JAFDED01 GB GCA 019310535.1
DSWW01
Syntrophia
JdFR-97
DAOWOC01 GB GCA 030642225.1
DSM-4660
Desulfobacteria
DTXE01 GB GCA 012962595.1
Desulfofervidia
Syntrophobacteria
QYQD01
Desulfobulbia
Dissulfuribacteria
Thermodesulfobacteria
Desulfobaccia
RBG-13-43-22
Desulfarculia
JACQYL01
JADFXK01
S145-22
Anaeroferrophillalia
Zymogenia
o__Desulfomonilales
g__B14-G9
g__JAGVNP01 GB GCA 020053185.1
g JAGGYK01
g__DATCTV01
g__JAKQBR01
g__JAYEKR01 GB GCA 034661115.1
g__JAFGIF01 GB GCA 016929755.1
g__DSBG01 GB GCA 011046135.1
g__MWEI01
g__DSAD01
GB GCA 002448235.1
GB GCA 029547185.1
GB GCA 035352355.1
GB GCA 022703355.1
GB GCA 002316295.1
SV3_maxbin2_001
GB GCA 001896555.1
Candidatus Phosphitivorax Phox-21
GB GCA 023230665.1
GB GCA 018436855.1
GB GCA 018433775.1
GB GCA 018433085.1
SV3_maxbin2_002
DPO139_4 maxbin2
DPO139_3 maxbin2_001
DPO139_2 concoct_28
DPO139_1 maxbin2_001
Ca. Phosphitivorax UBA1062
UBA1062
Desulfomonilia

**Supplemental Figure 9:** Phylogenomic tree of Phox-21 and representative bacterial genomes from the phylum Desulfobacterota from the GTDB. Clades are collapsed at the class level. The orange highlighted region indicates the class Desulfomonilia. The dark pink highlighted region represents the order UBA1062. The purple highlighted region represents the family and genus UBA1062. Nodes labeled "DPO139" are Phox-21 metagenome-assembled genomes (MAGs) generated in this study. Nodes labeled "SV3" are Phox-21 MAGs generated by Ewens et al. 2021. The node labeled "Candidatus Phosphitivorax Phox-21" is a MAG generated by Figueroa et al. 2017. Internal nodes with bootstrap values of >90% are indicated by purple circles.

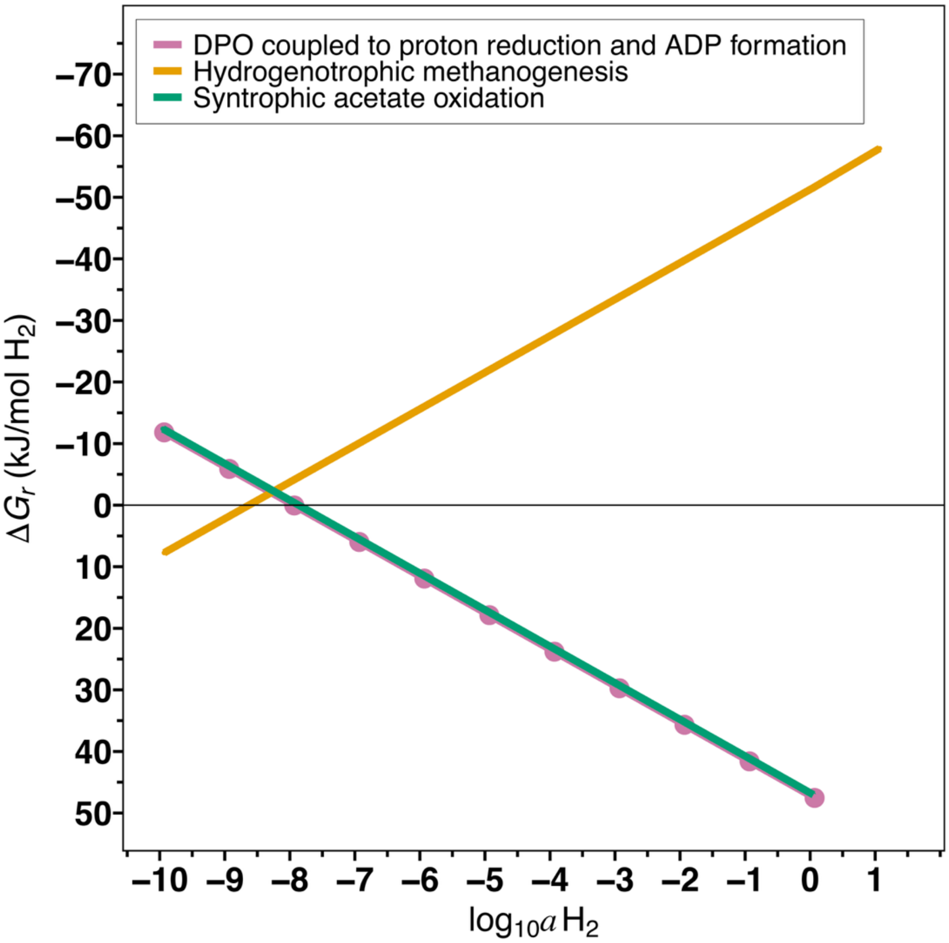


**Supplemental Figure 10:** Gibbs energies, $\Delta G_r$, of reactions 7, 9, and 18 (Table 1) as a function of hydrogen activity under simulated marine sediment conditions (4 ºC). Activities (Supp. Table 4) for products and reactants were calculated from the concentrations listed in Supp. Table 3. $\Delta G_r$ was calculated using activities determined from the following concentrations: 1 µM phosphite, 1µM acetate, and 100 µM methane, at 4 ºC.
